# CAR-engineered lymphocyte persistence is governed by a FAS ligand/FAS auto-regulatory circuit

**DOI:** 10.1101/2024.02.26.582108

**Authors:** Fei Yi, Tal Cohen, Natalie Zimmerman, Friederike Dündar, Paul Zumbo, Razan Eltilib, Erica J. Brophy, Hannah Arkin, Judith Feucht, Michael V. Gormally, Christopher S. Hackett, Korbinian N. Kropp, Inaki Etxeberria, Smita S. Chandran, Jae H. Park, Katharine C. Hsu, Michel Sadelain, Doron Betel, Christopher A. Klebanoff

## Abstract

Chimeric antigen receptor (CAR)-engineered T and NK cells can cause durable remission of B-cell malignancies; however, limited persistence restrains the full potential of these therapies in many patients. The FAS ligand (FAS-L)/FAS pathway governs naturally-occurring lymphocyte homeostasis, yet knowledge of which cells express FAS-L in patients and whether these sources compromise CAR persistence remains incomplete. Here, we constructed a single-cell atlas of diverse cancer types to identify cellular subsets expressing *FASLG*, the gene encoding FAS-L. We discovered that *FASLG* is limited primarily to endogenous T cells, NK cells, and CAR-T cells while tumor and stromal cells express minimal *FASLG*. To establish whether CAR-T/NK cell survival is regulated through FAS-L, we performed competitive fitness assays using lymphocytes modified with or without a FAS dominant negative receptor (ΔFAS). Following adoptive transfer, ΔFAS-expressing CAR-T and CAR-NK cells became enriched across multiple tissues, a phenomenon that mechanistically was reverted through *FASLG* knockout. By contrast, *FASLG* was dispensable for CAR-mediated tumor killing. In multiple models, ΔFAS co-expression by CAR-T and CAR-NK enhanced antitumor efficacy compared with CAR cells alone. Together, these findings reveal that CAR-engineered lymphocyte persistence is governed by a FAS-L/FAS auto-regulatory circuit.

## Introduction

Chimeric antigen receptor (CAR) T cell therapies have revolutionized the treatment of many B-cell malignancies^1–10^ and are now showing early signs of efficacy in solid cancers^11–17^. Despite ongoing progress, many patients who receive CAR therapies either fail to respond or develop acquired resistance, highlighting a critical need to further optimize current treatments. Multiple factors may contribute to disease progression following adoptive cell transfer (ACT). Broadly, these can be categorized as tumor-intrinsic, tumor microenvironmental, or CAR lymphocyte-intrinsic resistance mechanisms^18,19^. Among these, factors associated with the intrinsic properties of CAR-expressing lymphocytes are of particular interest because they are potentially modifiable during *ex vivo* cell manufacturing. Across disease types and CAR designs, the expansion and/or persistence of transferred cells is among the most consistent correlations associated with superior patient outcomes^2,3,7,8,20–22^. Emerging clinical data indicates this attribute holds true not only for CAR-T but also CAR-modified NK cells (CAR-NK)^23^, a lymphocyte subset with desirable features for allogeneic applications^24^. Thus, enhancing the survivability and durability of CAR-T and CAR-NK cells remains a major goal in the cell therapy field.

Unbiased genome-wide^25,26^ and focused^27–29^ CRISPR screens have recently identified *FAS* as a major determinant of antitumor T cell persistence under chronic antigen stimulation conditions. FAS is one of five tumor necrosis factor (TNF) superfamily death receptors that induces caspase-dependent apoptosis following engagement with an extracellular ligand^30,31^. These findings provide a strong rationale to develop strategies that disable FAS-signaling in receptor engineered T cells^27,28,32–35^. However, while the role of FAS in regulating naturally occurring T cell homeostasis is well established, whether this pathway governs CAR-NK longevity remains unknown. Moreover, the source of FAS ligand (FAS-L) in cancer patients and whether the benefit of FAS antagonism occurs only when the ligand is expressed by specific cell types remains incompletely understood.

In the present study, we sought to address three critical gaps in knowledge. First, we sought to define which cell types express *FASLG* in cancer patients. Second, we sought to establish whether CAR-engineered lymphocyte persistence is negatively self-regulated by *FASLG*. Finally, we sought to determine whether *FASLG* is required for on-target CAR-T and CAR-NK effector functions against B-cell malignancies. Our findings indicate that strategies which disrupt FAS-signaling are broadly applicable across cellular immunotherapy platforms and agnostic of cancer type and lymphocyte subset transferred.

## Results

### FASLG is expressed by endogenous and CAR-expressing lymphocytes

We previously established that ∼73% of human tumor types represented in The Cancer Genome Atlas overexpress *FASLG*, the gene encoding FAS ligand (FAS-L), compared with matched normal tissues^32^. Because this analysis was performed using bulk RNA sequencing, the identity of specific ligand-expressing cell type(s) could not be unambiguously resolved. To precisely define which cellular subsets express *FASLG*, we generated an integrated human single-cell transcriptomic atlas comprised of 244,809 immune and non-immune cells using publicly available datasets **(Extended Data Fig. 1)**. To capture phenotypic heterogeneity, cells were interrogated from 37 patients with diverse hematologic and solid cancers and four healthy donors. After applying stringent quality controls and a standardized analysis pipeline, uniform manifold approximation and projection (UMAP) visualization revealed nine distinct clusters (**Fig. 1a**). Expression of canonical marker genes identified 18 major cell types that were subcategorized as follows: malignant, stromal, non-lymphoid immune, and lymphocyte subsets.

**Fig. 1:**
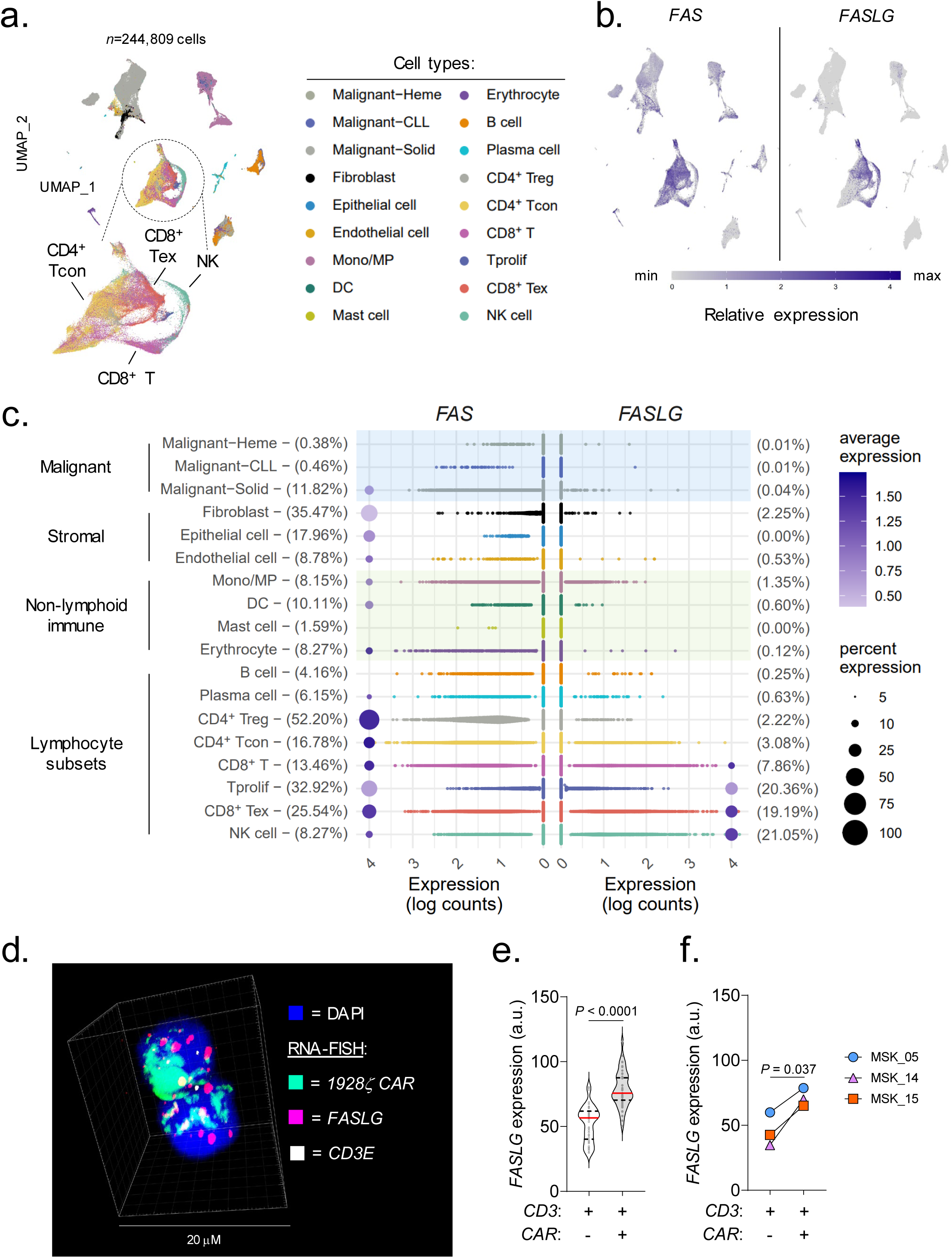
*FASLG* is expressed by endogenous and CAR-expressing lymphocytes. (**a**) UMAP visualization of scRNA-seq from *n*=244,809 immune and non-immune cells obtained from the tumor and peripheral blood of patients with *n*=10 hematologic cancers, *n*=27 solid cancers, and peripheral blood from *n*=4 healthy donors. Each dot represents an individual cell assigned to one of 18 inferred cell types. (**b**) Log-transformed normalized gene expression values for *FAS* and *FASLG* overlaid on the UMAP coordinates defined in panel (**a**). (**c**) Comparison of the frequency and magnitude of *FAS* and *FASLG* expression by individual cells assigned to each inferred cell type identified in the UMAP visualization. Bubble size represents the frequency of each cell type that expresses the indicated gene and color indicates the relative intensity of expression. (**d**) Representative immunofluorescent confocal image, (**e**) summary violin plots, and (**f**) patient-level analyses quantifying *FASLG* mRNA expression by endogenous and CAR-expressing T cells in the bone marrow of patients with B-ALL treated with a 1928ζ CAR. Samples were co-hybridized with DAPI (blue) and multiplexed RNA-FISH probes specific for the mRNA sequence of the CAR’s single-chain variable fragment (scFv) (green), *CD3E* mRNA (white), and *FASLG* mRNA (red). Data shown is derived from *n*=3 patients. Violin distributions are centered around the median (red horizontal line) with quartiles ranges displayed above and below (dashed horizontal lines). The maxima and minima are represented by the top and bottom of each plot. Each dot represents mean *FASLG* mRNA expression within a particular cell type from an annotated region of interest. *P*-values calculated using a two-sided Student’s t-test. a.u. = arbitrary fluorescence units.

Overlay of mRNA expression for specific genes onto the UMAP coordinates revealed that *FAS* was broadly distributed across many cell types (**Fig. 1b**). This includes a subset of normal and cancer-associated fibroblasts, dendritic cells, T regulatory cells, proliferating T cells, and exhausted T cells (**Fig. 1c**), findings consistent with prior studies^36–39^. By contrast, we discovered that *FASLG* expression was highly restricted and limited primarily to T and NK cells. In this analysis, there was minimal to no *FASLG* expression detected within malignant and stromal cells.

Having resolved the landscape of *FASLG* expression by endogenous cells, we next sought to measure expression of this ligand by CAR-engineered T cells following adoptive transfer into patients. To accomplish this, we developed and validated a multiplexed fluorescent RNA *in situ* hybridization (RNA-FISH) assay to quantify co-expression of mRNA for a 1928ζ CAR transgene, *CD3E*, and *FASLG* at single-cell resolution (**Extended Data Fig. 2**). Using bone marrow (BM) biopsy samples from patients with B-ALL treated in the context of a clinical trial (NCT01044069)^20^, we found that CAR^+^ T cells consistently expressed significantly higher levels of *FASLG* compared with endogenous T cells (**Fig. 1d-f**). Taken together, we conclude that *FASLG* expression within cancer patients is highly restricted and primarily associated with a subset of endogenous lymphocytes and CAR-modified T cells.

### CAR-T derived FASLG auto-regulates cellular persistence in vivo

Nearly all commercial and experimental CAR-engineered cell products are administered following a lymphocyte-depleting chemotherapy regimen^40^. Consequently, CAR-modified cells represent a significant proportion of the total circulating lymphocyte pool in the weeks following adoptive transfer^21,41–43^. We hypothesized that activation-dependent FAS-L expression by CAR-T cells will engage FAS^+^CAR-T cells to induce apoptosis and limit cellular persistence. To test this, we performed a competitive fitness assay using two phenotypically discernable populations of human CAR-T cells that are either responsive or unresponsive to FAS-signaling (**Fig. 2a**). In this experiment, T cells were individually transduced with one of two multi-cistronic vectors: *1*) a vector encoding a 1928ζ CAR, a FAS dominant negative receptor (ΔFAS) with a truncation in the death domain, and truncated epidermal growth factor receptor (tEGFR), or *2*) a vector encoding an identical CAR and truncated low-affinity neuronal growth factor (tLNGFR). tEGFR and tLNGFR are functionally inert cell surface molecules that are coordinately expressed with the CAR transgene^44,45^, enabling mixed populations of CAR-T cells to be tracked longitudinally. We confirmed that T cell expression of ΔFAS blocks FAS ligand-induced apoptosis but has no detrimental impact on antigen-dependent CAR-T cytokine release or cytolytic functions *in vitro* (**Extended Data Fig. 3**). tEGFR^+^ and tLNGFR^+^ T cells were combined in a ∼1:1 ratio and adoptively co-transferred into NOD/SCID/γc^-/-^ (NSG) mice bearing established Nalm6 leukemia. At the time of transfer, tEGFR^+^ and tLNGFR^+^ T cells were comprised of comparable frequencies of T stem cell memory (T_SCM_) and T central memory (T_CM_) cells, subsets associated with superior CAR-T persistence (**Fig. 2b**)^21,22,46^. After one month, we measured the proportion of FAS-signaling competent and incompetent CAR-T cells in multiple tissues, including the blood, BM, liver, and spleen. We consistently observed significant enrichment for ΔFAS-expressing T cells across all sites with the greatest magnitude of skewing occurring in the BM, a dominant site of disease in this model (**Fig. 2c,d**). These results demonstrate that FAS-signaling regulates the survival of CAR-T cells within a tumor-bearing host in a cell-intrinsic manner.

**Fig. 2:**
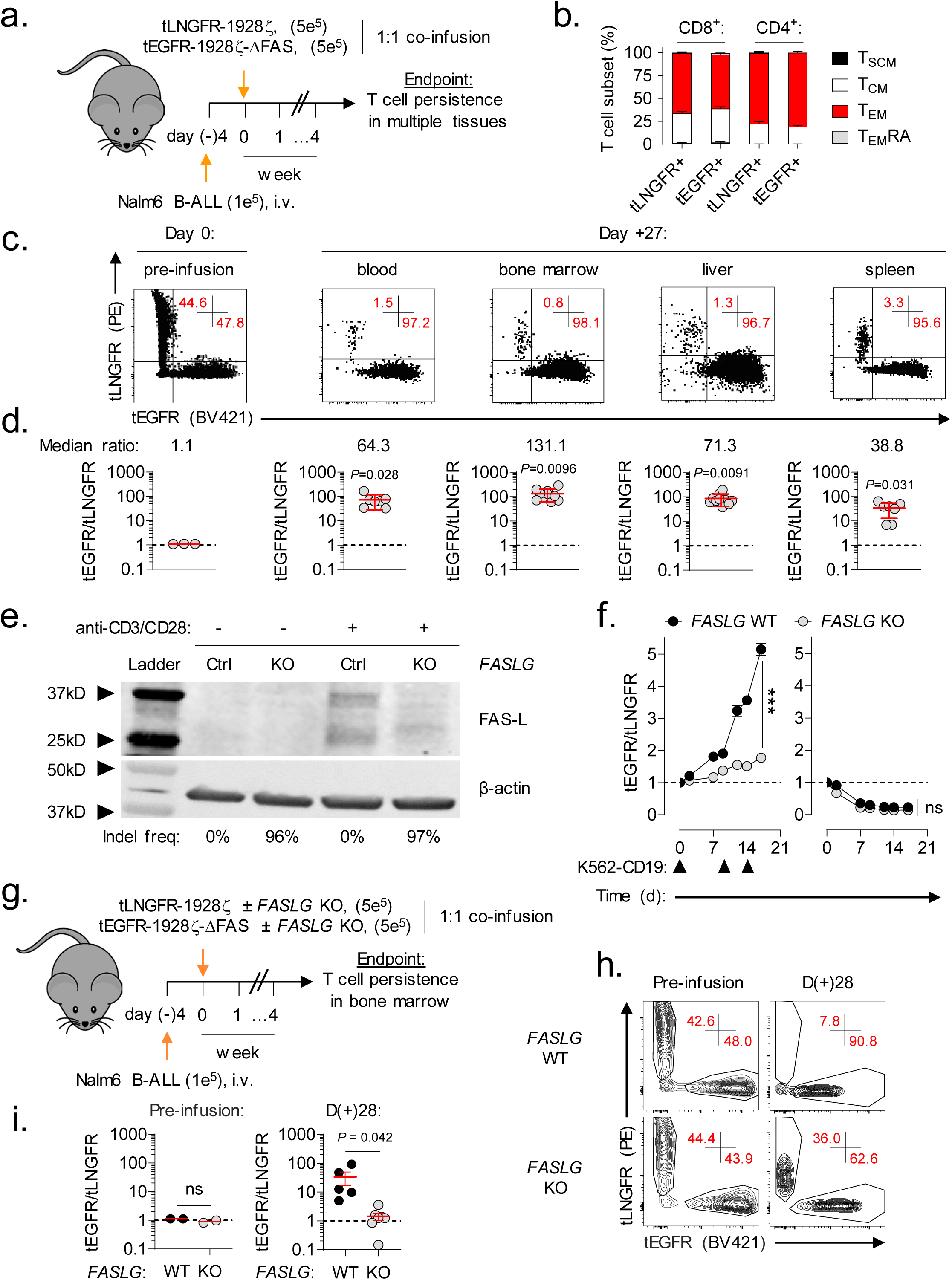
CAR-T derived *FASLG* auto-regulates cellular persistence *in vivo*. (**a**) Schematic overview of the experimental design to test the *in vivo* persistence of human T cells that express a 1928ζ CAR ± a FAS dominant negative receptor (ΔFAS) in tumor-bearing mice. T cells were co-transferred at a ∼1:1 ratio into NSG mice bearing established Nalm6 B-cell acute lymphoblastic leukemia (B-ALL) and tracked based on expression of tLNGFR or tEGFR. (**b**) Percentage of CD8^+^ and CD4^+^ T cells with a given memory phenotype at the time of adoptive transfer following transduction with tLNGFR-1928ζ or tEGFR-1928ζ-ΔFAS. Bar graphs displayed as mean ± s.e.m. using *n*=3 biological replicates. (**c**) Representative FACS and (**d**) summary scatter plots measuring the ratio of tEGFR^+^ to tLNGFR^+^ T cells at the time of infusion and four weeks following adoptive transfer. *P*-values calculated based on comparison to the infusion product using a two-sided Student’s t-test. (**e**) Western blot for FAS-L protein expression in lysates from control or FASLG-KO 1928ζ CAR-transduced T cells. Cells were analyzed at rest and 48h after anti-CD3/CD28 restimulation. The frequency of frameshift Indels in *FASLG* for each cell type are shown beneath each lane. (**f**) Relative antigen-driven in vitro expansion of control and *FASLG*-KO 1928ζ CAR-T cells with or without ΔFAS co-expression. CAR-T cells were combined in ∼1:1 ratio on day 0 and serially restimulated at indicated time points (▴) with K562-CD19 *FASLG*-KO leukemia cells (left panel) or left unstimulated as controls (right panel). Data is displayed as the mean ratio of tEGFR/tLNGFR T cells ± s.e.m. using *n*=3 biological replicates. Groups were compared using a paired two-tailed Student’s T test for accumulated differences between each time point. (**g**) Schematic overview of the experimental design to test the influence of CAR-T derived *FASLG* on *in vivo* persistence in mice bearing established Nalm6 B-ALL. Control or *FASLG*-KO tLNGFR-1928ζ CAR-T cells were co-transferred at a ∼1:1 ratio with control or *FASLG*-KO tEGFR-1928ζ-ΔFAS CAR-T cells into Nalm6 B-ALL bearing NSG mice. (**h**) Representative FACS and (**i**) summary scatter plot comparing the ratio of tEGFR to tLNGFR cells at the time of infusion and four weeks following adoptive transfer. Symbols displayed as mean ± s.e.m. Groups compared using a two-sided Student’s T test. ns, not significant (*P*>0.05).

*FASLG* expression is primarily restricted to T cells and NK cells (**Fig. 1b**) and adoptively transferred T cells remain the dominant lymphocyte population following adoptive transfer. We therefore next sought to establish whether CAR-T derived *FASLG* is sufficient to drive population skewing. To address this question, we ablated *FASLG* in CAR-T cells using Cas9 ribonucleoprotein (RNP)-mediated gene knock out (KO)^47^. Tracking of insertions/deletions by decomposition (TIDE) analysis^48^ confirmed a high frequency (>96%) of frameshift mutations at the *FASLG* locus in *FASLG*-KO CAR-T cells and no measurable disruption in control-edited cells. To test the activation-dependence of FAS-L protein expression by CAR-T cells, we performed Western blot on cell lysates from control and *FASLG*-KO T cells at rest and 48h after anti-CD3/CD28 restimulation. Minimal to no FAS-L protein was measured in resting CAR-T cells (**Fig. 2e**). After restimulation, two molecular weight bands measuring ∼37 kD and ∼26 kD were detected in control-edited cells. These correspond to the previously described integral membrane and metalloproteinase-generated soluble forms of FAS ligand, respectively^49^. The presence of both bands was nearly completely abrogated from activated *FASLG*-KO cells, confirming gene disruption.

Having established the activation-dependence of FAS-L protein expression by CAR-T cells, we next tested the role of CAR-T derived *FASLG* on cellular persistence following antigen encounter. First, we measured the influence of repetitive *in vitro* stimulation using CD19^+^K562 *FASLG*-KO leukemia cells on the distribution of ΔFAS to wild-type (WT) FAS-expressing CAR-T cells. In this experiment, ΔFAS/tEGFR-expressing CAR-T cells were added in a 1:1 ratio with tLNGFR-expressing CAR-T cells. The mixed population was serially activated through the addition of CD19^+^K562 cells on days zero, nine, and 14 or left untreated as controls. Beginning with the first round of tumor stimulation, the ratio of tEGFR/tLNGFR cells became progressively enriched for FAS-signaling deficient CAR-T cells (**Fig. 2f**, left). Knock out of *FASLG* significantly reduced the magnitude of CAR-T cell population skewing such that the ratio of the two cell types remained close to one. In the absence of stimulation, there was no significant difference between *FASLG* intact and KO cells and the ratio of tEGFR/tLNGFR declined with time (**Fig. 2f**, right). This indicates that disruption of FAS-signaling in human CAR-T cells does not lead to potential safety considerations, such as uncontrolled, antigen-independent cell accumulation.

Based on these *in vitro* findings, we sought to establish whether CAR-T derived *FASLG* is sufficient to control cellular persistence *in vivo*. To test this, tEGFR-1928ζ-ΔFAS and tLNGFR-1928ζ expressing T cells were recombined in a 1:1 fashion after *FASLG* or control-KO (**Fig. 2g**). The mixed T cell populations were co-infused into NSG mice bearing established Nalm6 tumors and the ratio of T cell subsets was measured in BM one month after transfer. Consistent with our prior results, the balance of control-edited T cells became altered in favor of ΔFAS-expressing CAR-T cells. By contrast, the median ratio of the two T cell populations remained close to one with *FASLG*-KO (**Fig. 2h,i**). Thus, we conclude that activation-induced CAR-T FAS-L expression restrains the persistence of adoptively transferred CAR-T cells in a FAS-dependent manner.

### Disabling FAS-signaling enhances CAR-T antitumor efficacy in vivo

Based on the finding that FAS-L regulates the persistence of CAR-T cells, we postulated that T cell-intrinsic disruption of FAS-signaling would enhance CAR-dependent antitumor efficacy *in vivo*. To test this hypothesis, we transduced human T cells with vectors encoding tEGFR/1928ζ with or without ΔFAS. As a control for antigen-specificity, an aliquot of T cells from the same donor were transduced with tEGFR alone. Transduced T cells were adoptively transferred into NSG mice bearing established luciferase-expressing Nalm6 tumors at a CAR-T cell dose previously determined to be non-curative (**Fig. 3a**)^50,51^. Changes in tumor burden, assessed using bioluminescence imaging (BLI), and overall animal survival were measured. Compared with tEGFR^+^ control cells, we found that transfer of either CAR-expressing T cell population delayed tumor progression and significantly improved overall survival (*P*<0.0001) (**Fig. 3b,c**). Consistent with the enhanced persistence potential of human CAR-T cells which co-express ΔFAS, mice treated with FAS-signaling disrupted CAR-T cells experienced prolonged tumor control. This resulted in significantly improved overall survival (*P*=0.0023).

**Fig. 3:**
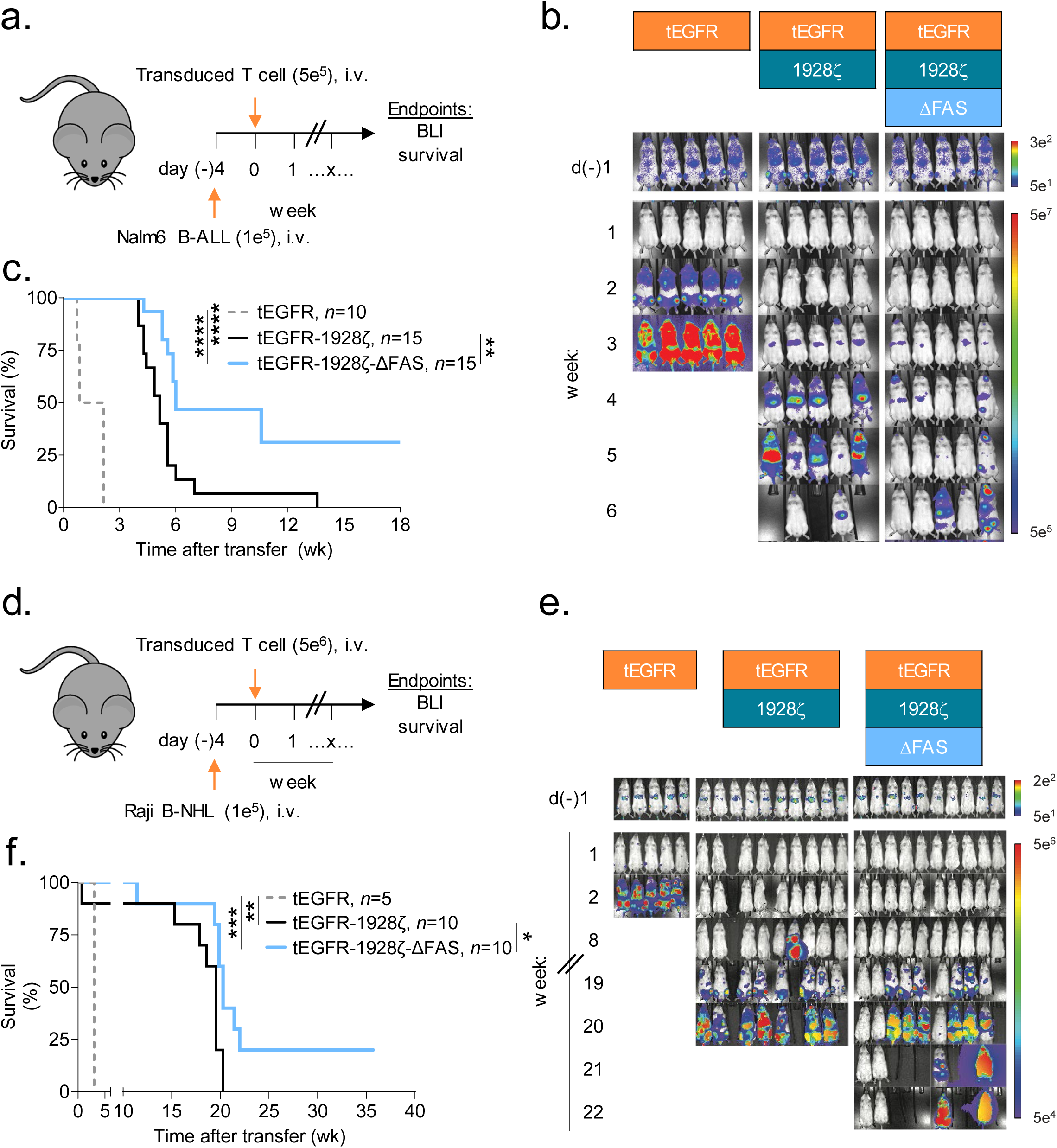
Disabling FAS-signaling enhances CAR-T antitumor efficacy *in vivo*. (**a**) Experimental design to compare the *in vivo* antitumor efficacy of human T cells that express a 1928ζ CAR ± a FAS dominant negative receptor (ΔFAS) against established Nalm6-luciferase (Luc) B-cell acute lymphoblastic leukemia (B-ALL). (**b**) Bioluminescence imaging (BLI) and (**c**) survival curves for Nalm6 B-ALL bearing NSG mice treated by i.v. injection with 5 x 10^5^ tEGFR^+^ T cells transduced with the indicated vectors. Pooled survival data from two identically performed experiments is shown in (**c**) plotted as a Kaplan–Meier survival curve (tEGFR alone, *n* = 10; tEGFR-1928ζ, *n* = 15; tEGFR-1928ζ-ΔFAS, *n* = 15). \*\*\*\**P*<0.0001 and \*\**P*=0.0023 using a log-rank test. (**d**) Experimental design to compare the in vivo antitumor efficacy of human T cells that express a 1928ζ CAR ± ΔFAS against established Raji-Luc B-cell non-Hodgkin’s lymphoma (B-NHL). (**e**) Bioluminescence imaging and (**f**) survival curves for mice bearing Raji B-NHL and treated by i.v. injection with 5 x 10^6^ tEGFR^+^ T cells transduced with the indicated vectors. Survival data is plotted as a Kaplan–Meier survival curve (tEGFR alone, *n* = 5; tEGFR-1928ζ, *n* = 10; tEGFR-1928ζ-ΔFAS, *n* = 10). \*\*\**P*=0.0002, \*\**P*=0.0043, and \**P*=0.0169 using log-rank test. wk, week.

We next investigated the generalizability of employing cell autonomous FAS-signaling blockade to enhance the potency of CAR-T cells using a second B-cell malignancy. NSG mice were pre-implanted with luciferase-expressing Raji cells, a model for aggressive B-cell non-Hodgkin lymphoma (B-NHL). Tumor-bearing animals were treated with a single i.v. infusion of tEGFR, tEGFR/1928ζ, or tEGFR/1928ζ/ΔFAS transduced T cells (**Fig. 3d**). Groups that received CAR-expressing T cells controlled tumor growth and had significantly improved overall survival compared with the tEGFR only controls (*P*≤0.0043) (**Fig. 3e,f**). Similar to results in the pre-B ALL model, mice that received ΔFAS co-expressing CAR-T cells had significantly enhanced overall survival compared with mice receiving CAR-T cells alone (*P*=0.0169). Taken together, we conclude that cell-intrinsic blockade of FAS-signaling significantly enhances CAR-mediated tumor control of CD19-expressing malignancies *in vivo*.

### FAS-L/FAS-signaling is dispensable for CAR-T antitumor efficacy

Having established that *FASLG* expression by CAR-T cells negatively regulates cellular persistence, we next sought to define the role of this molecule in CAR-mediated tumor lysis. First, we compared the *in vitro* antitumor potency of *FASLG*-KO versus control-edited 1928ζ CAR-T cells using an IncuCyte live-imaging cytotoxicity assay (**Fig. 4a**). We^52^ and others^35,53^ previously determined that human effector memory T cells are differentially sensitive to FAS-mediated apoptosis relative to naïve (T_N_)-derived T cells. To avoid confounding based on differences in T-cell differentiation at the start of the tumor co-culture, we enriched for T_N_ prior to CAR transduction in these experiments. At high effector to target (E:T) ratios against Nalm6 (1:2 and 1:4), we observed no measurable differences in tumor lysis efficiencies between *FASLG*-KO and control-edited T cells (**Fig. 4b**). At lower E:T ratios (≥1:8), however, we observed significantly greater CAR-mediated cytotoxicity by the *FASLG*-KO groups. In the absence of CAR-T cells, we found that a recombinant version of FAS-L oligomerized through a leucine zipper domain (lzFAS-L)^52,54^ similarly had a minimal impact on the growth kinetics of Nalm6 and Raji cells (**Fig. 4c**). By contrast, lzFAS-L exposure caused a dose-dependent depletion of activated T cells.

**Fig. 4:**
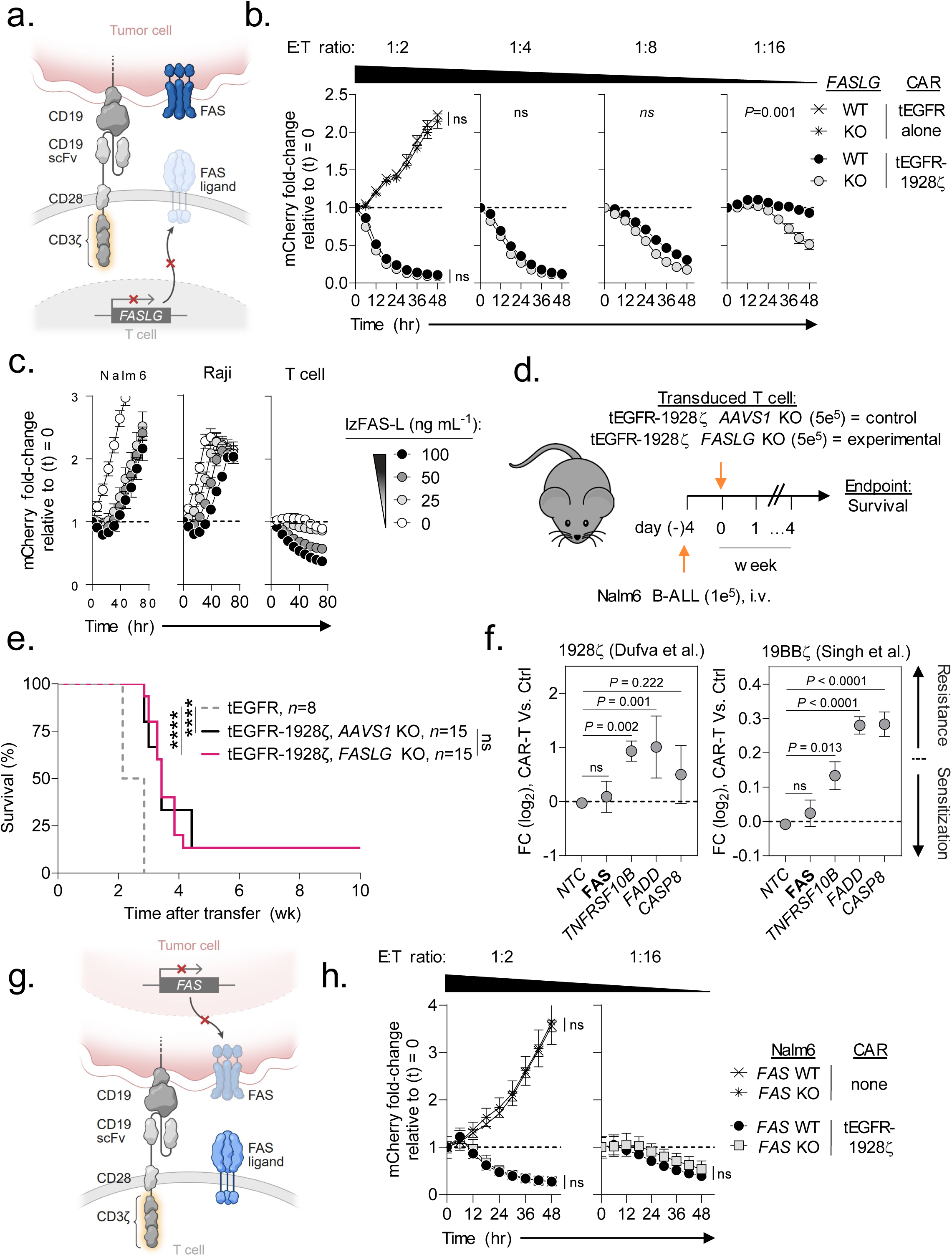
FAS-L/FAS-signaling is dispensable for CAR-T antitumor efficacy. (**a**) Schematic for the CRISPR/Cas9-mediated knockout (KO) of *FASLG* (experimental) or *AAVS1* (control) in human CD8^+^ T cells expressing a 1928ζ CAR. (**b**) Cytolytic activity of *FASLG*-KO versus *AAVS1*-KO 1928ζ CAR-T cells against Nalm6/mCherry at indicated effector to target (E:T) ratios. Data shown as mean ± s.e.m. using *n*=3 biological replicates per condition. Statistical comparisons performed using a one-way ANOVA. ns = not significant (*P*>0.05). (**c**) Growth kinetics of Nalm-6 B-ALL, Raji B-NHL, or activated T cells in the presence or absence of lzFAS-L. Each cell type was transduced with mCherry. Data shown as mean ± s.e.m. using *n*=3 biological replicates per condition following incubation with an indicated concentration of lzFAS-L. hr, hour. (**d**) Experimental design and (**e**) Kaplan–Meier survival curve comparing the *in vivo* antitumor efficacy of human CD8^+^ T cells that express a 1928ζ CAR with *FASLG*-KO or *AAVS1*-KO against established Nalm-6 B-ALL. tEGFR alone, *n*=8; tEGFR-1928ζ *AAVS1*-KO, *n*=15; tEGFR-1928ζ *FASLG*-KO, *n*=15. \*\*\*\**P*< 0.0001 and ns (not significant, *P*>0.05) using log-rank test. wk, week. (**f**) Scatter plots displaying the enrichment or depletion of synthetic guide RNAs (sgRNAs) targeting indicated genes in the death receptor pathway by Cas9-expressing Nalm-6 B-ALL cells. Tumor cells were placed under selection by T cells transduced with a 1928ζ CAR (left), a 41BBζ CAR (right), or left non-transduced as specificity controls (Ctrl). Data re-analyzed from two published genome-scale CRISPR/Cas9 screens^56,57^ and is shown as mean log_2_ fold-change (FC) ± s.e.m of sgRNAs targeting indicated genes. NTC = non-targeted control sgRNAs. Gene level significance was determined using a one-way ANOVA corrected for multiple comparisons by Dunnett’s test. (**g**) Schematic for the CRISPR/Cas9-mediated knockout (KO) of FAS using an individual sgRNA in Nalm6 B-ALL. (**h**) Time-dependent cytolytic activity of 1928ζ CAR-T cells against FAS-KO versus FAS-wild type (WT) Nalm6/mCherry cells at a high (left) or low (right) E:T ratio. Data shown as mean ± s.e.m. using *n*=3 biological replicates per condition and time point. Statistical comparisons were performed using a one-way ANOVA.

We next sought to establish whether *FASLG* was dispensable for on-target CAR-T mediated antitumor efficacy *in vivo*. To test this, tEGFR/1928ζ-transduced T cells underwent Cas9 RNP-mediated editing of either *FASLG* or adeno-associated virus site 1 (*AAVS1*)^55^ as a control (**Fig. 4d**). Gene edited CAR-T cells or T cells transduced with tEGFR alone were then transferred into mice bearing established Nalm6 tumors. Relative to tEGFR alone, both *FASLG*-KO and *AAVS1*-KO CAR-T cells significantly enhanced overall animal survival (**Fig. 4e**). Similar to our *in vitro* findings, there was no detriment in *in vivo* antitumor efficacy by *FASLG*-KO CAR-T cells.

To provide additional evidence that CAR-mediated cancer cell lysis occurs independently of the FAS-L/FAS pathway, we re-analyzed results from two published genome-scale CRISPR screens^56,57^. These screens placed Nalm6 cells under selection by T cells expressing either a 1928ζ or 41BB (BB)-containing (19BBζ) CAR to identify tumor-intrinsic resistance mechanisms. In neither screen was enrichment for tumor clones expressing synthetic guide RNAs (sgRNAs) targeting *FAS* observed relative to non-targeted control sgRNAs (**Fig. 4f**). We validated this finding using a unique *FAS*-targeting sgRNA sequence not contained in either screen and 1928ζ CAR-T cells at high (1:2) and low (1:16) E:T ratios (**Fig. 4g**,**h**). Unlike *FAS*, significant enrichment for sgRNAs targeting *TNFRSF10B*, the gene encoding the alternative death receptor TRAIL-R2, was observed using either CAR design indicating therapeutic resistance. Moreover, sgRNAs targeting downstream mediators of TRAIL-R2-signaling, including *FADD* and *CASP8*, were also enriched in both screens. Consistent with these findings, we discovered that KO of the ligand for TRAIL-R2 (*TNFSF10*, also known as *TRAIL*) in 1928ζ CAR-T cells significantly impaired Nalm6 lysis (**Extended Data Fig. 4**). Taken together, we conclude that on-target CAR-mediated antitumor efficacy against B-cell malignancies can occur independent of the FAS-L/FAS pathway.

### CAR-NK survival is regulated by a FAS/FAS ligand auto-regulatory circuit

Results from our single-cell transcriptomic atlas revealed that in addition to T cells, a subset of human NK cells also express high levels of *FAS* and *FASLG*. Based on this finding, we next sought to investigate whether naturally occurring and CAR-engineered NK cells express FAS protein and are sensitive to FAS-L induced apoptosis. Resting cord-blood derived NK cells displayed minimal FAS protein; following activation, however, expression was significantly up-regulated (**Fig. 5a,b**). We next tested whether 1928ζ CAR-transduced NK cells and activated but non-transduced NK cells are responsive to FAS-L. In the absence of exogenous FAS-L, non-transduced and CAR-transduced NK cells expressed minimal to no activated caspase 3/7 and annexin V, markers of early and late apoptosis (**Fig. 5c**). Expression of both markers significantly increased following exposure to lzFAS-L, a process that could be blocked by co-expression of ΔFAS.

**Fig. 5:**
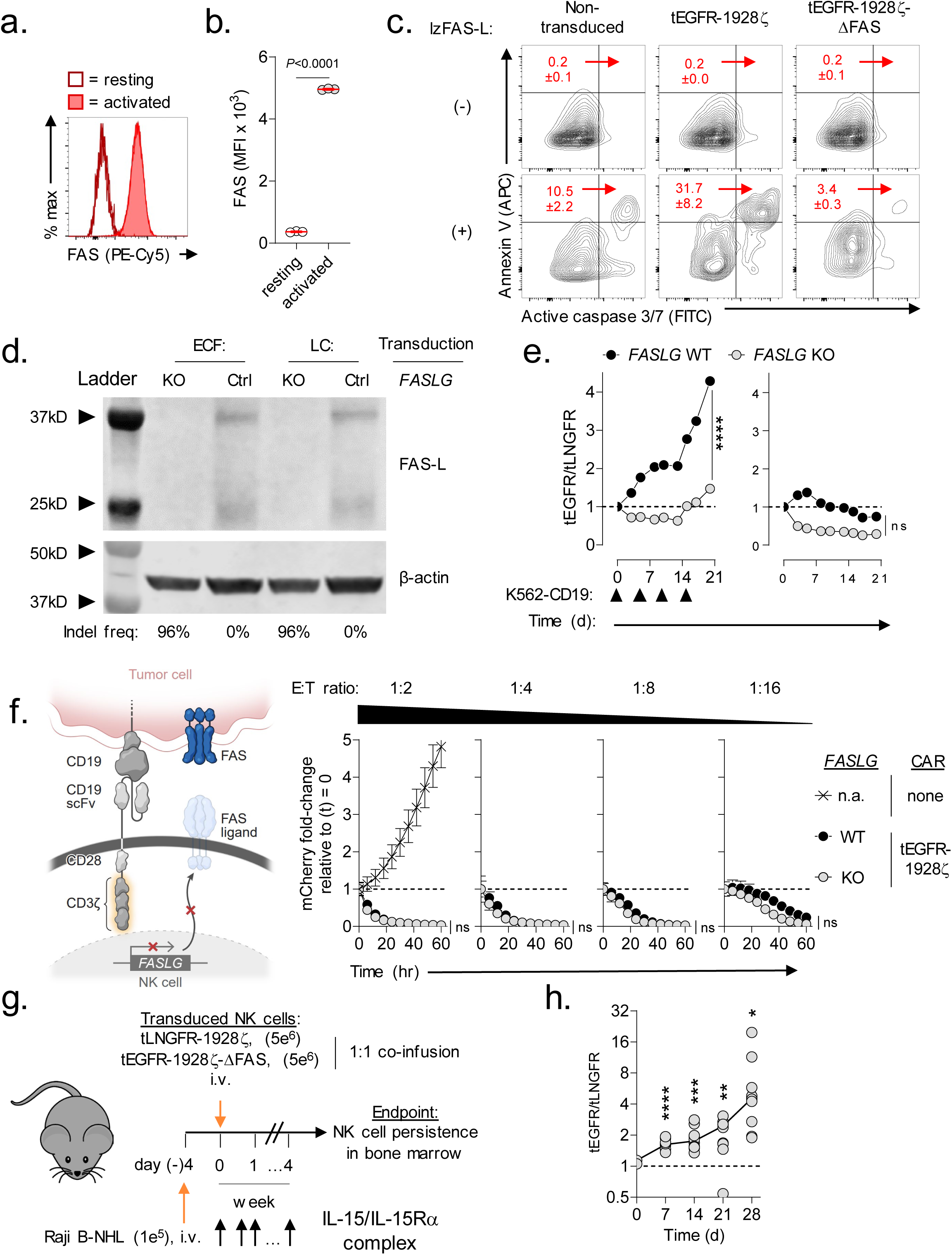
CAR-NK survival is regulated by a FAS/FAS ligand auto-regulatory circuit. (**a**) Representative FACS and (**b**) summary scatter plot quantifying FAS expression on cord blood-derived human NK cells at rest and 5d following activation with irradiated K562 Clone 9 cells. Data shown as mean ± s.e.m. using *n*=3 biological replicates per condition. Statistical analysis performed by two-sided Student’s T test. (**c**) Representative FACS plots quantifying lzFAS-L induced apoptosis in activated non-transduced NK cells or NK cells transduced with tEGFR-1928ζ or tEGFR-1928ζ-ΔFAS. Numbers indicate the mean ± s.e.m. of activated caspase 3/7+/annexin V^+^ cells for each condition. (**d**) Western blot for FAS-L protein expression in lysates from *FASLG*-KO or control (Ctrl) NK cells transduced with tEGFR-1928ζ-ΔFAS (ECF) or tLNGFR-1928ζ (LC). The frequency of frameshift Indels in *FASLG* are shown beneath each lane. (**e**) Relative antigen-driven *in vitro* expansion of control and *FASLG*-KO 1928ζ CAR-NK cells with or without ΔFAS co-expression. CAR-T cells were combined in ∼1:1 ratio on day 0 and serially restimulated at indicated time points (▴) with K562-CD19 FASLG-KO cells (left panel) or left unstimulated as controls (right panel). Data displayed as the mean ratio of tEGFR^+^ to tLNGFR^+^ NK cells ± s.e.m. using *n*=3 biological replicates. Groups compared using a paired two-tailed Student’s T test for accumulated differences between each time point. \*\*\*\**P*<0.0001, ns = not significant (*P*>0.05). (**f**) Cytolytic activity of *FASLG*-KO versus *FASLG*-wild type (WT) tEGFR-1928ζ CAR-NK cells against Raji/mCherry at indicated effector to target (E:T) ratios. Data shown as mean ± s.e.m. using *n*=3 biological replicates per condition. Statistical comparisons performed using a one-way ANOVA. ns = not significant (*P*>0.05), n.a. = not applicable. (**g**) Schematic overview of the experimental design to test the in vivo persistence of NK cells that express a 1928ζ CAR ± a FAS dominant negative receptor (ΔFAS) in Raji B-NHL bearing mice. (**h**) Scatter plot comparing the ratio of tEGFR^+^ to tLNGFR^+^ CAR-NK cells at the time of infusion and at indicated time points following adoptive transfer in the bone marrow of Raji B-NHL bearing mice. *P*-values calculated based on comparison to the infusion product using an unpaired, two-sided, Welch’s t-test. \*\*\*\**P*<0.0001, \*\*\**P*=0.0006, \*\**P*=0.0026, \**P*=0.0173.

We next determined whether human CAR-transduced NK cells express FAS-L protein. Lysates from NK cells transduced with tEGFR/1928ζ/ΔFAS or tLNGFR/1928ζ were probed with an anti-FAS ligand antibody by Western blot. As a specificity control, an aliquot of NK cells transduced with each vector underwent CRISPR-mediated *FASLG* editing and a high Indel frequency (≥96%) at the gene’s locus was confirmed. Similar to findings using CAR-T cells, we identified two bands with molecular weights corresponding to the membrane bound and soluble forms of FAS-L (**Fig. 5d**). Both bands were nearly completely ablated following *FASLG* gene editing, confirming successful KO in NK cells.

We next asked whether a FAS-L/FAS circuit regulates the survival of CAR-NK cells. NK cells were individually transduced with either tLNGFR/1928ζ or tEGFR/1928ζ/ΔFAS. tLNGFR and tEGFR-expressing CAR-NK cells were recombined in a ∼1:1 ratio and the mixed populations were serially restimulated with CD19^+^K562 *FASLG*-KO cells or left unstimulated as controls. Following each round of tumor restimulation, we measured progressive accumulation of ΔFAS-expressing CAR-NK cells (**Fig. 5e, left**). Population skewing was FAS-L dependent as *FASLG*-KO caused the ratio of tEGFR/tLNGFR NK cells to remain close to one. In the absence of antigen restimulation, the proportion of each CAR-NK population remained stable over time indicating that *FASLG*-induced fratricide was activation dependent (**Fig. 5e, right**).

Naturally occurring NK cells can eliminate pathogen-infected and cancer cells through multiple mechanisms, including exocytosis of preformed cytotoxic granules and death receptor engagement^58^. To test whether CAR-NK mediated killing of a B-lymphoid malignancy is FAS-L dependent, we compared the cytolytic efficiency of CAR-transduced NK cells with or without *FASLG*-KO. To distinguish between innate versus CAR-dependent effector functions, we measured killing against Raji B-NHL across a range of E:T ratios. Similar to results using CAR-T cells, we found that *FASLG* expression was also dispensable for CAR-NK cytotoxicity (**Fig. 5f**).

Finally, we sought to measure whether cell-intrinsic disruption of FAS-signaling could enhance the *in vivo* persistence of CAR-NK cells within tumor-bearing hosts. ΔFAS/tEGFR and tLNGFR CAR-NK cells were recombined in a 1:1 fashion and the mixed population was co-infused into Raji B-NHL tumor-bearing NSG mice (**Fig. 5g**). Beginning the day of NK transfer, an extended half-life variant of IL-15 was administered by intraperitoneal (IP) injection twice weekly to model the physiologic effect of lymphodepletion^59^. The ratio of the two engineered NK cell populations was measured in the infusion product and serially over time in the BM of recipient mice. We found that CAR-NK which co-express ΔFAS had significantly enhanced persistence relative to CAR-NK cells alone, resulting in progressive skewing in the tEGFR/tLNGFR ratio (**Fig. 5h**). Taken together, we conclude that a FAS-L/FAS auto-regulatory circuit controls the persistence of CAR-NK cells.

### ΔFAS enhances the in vivo antitumor efficacy of ITAM calibrated CAR-NK cells

CD3ζ-containing CAR designs are functional when expressed by NK cells^60–62^. Similar to T cells, NK cells use adapter proteins containing immunoreceptor tyrosine-based activation motifs (ITAMs) to drive downstream signaling from cross-linked surface receptors^63^. Recently, we reported that an ITAM-calibrated 1928ζ CAR (henceforth 1XX CAR) enables superior T cell persistence and antitumor efficacy compared with a conventional 1928ζ CAR^64^. Whether the 1XX 1928ζ CAR also provides for superior antitumor functions when expressed by NK cells has previously not been tested. To address this question, we transduced NK cells with either the native or 1XX variants of the 1928ζ CAR and measured time-dependent Raji B-NHL cytolysis across a range of E:T ratios (**Fig. 6a**). Although both groups of CAR-NK cells performed similarly at high E:T ratios, NK cells which expressed the 1XX CAR were superior at lower E:T ratios.

**Fig. 6:**
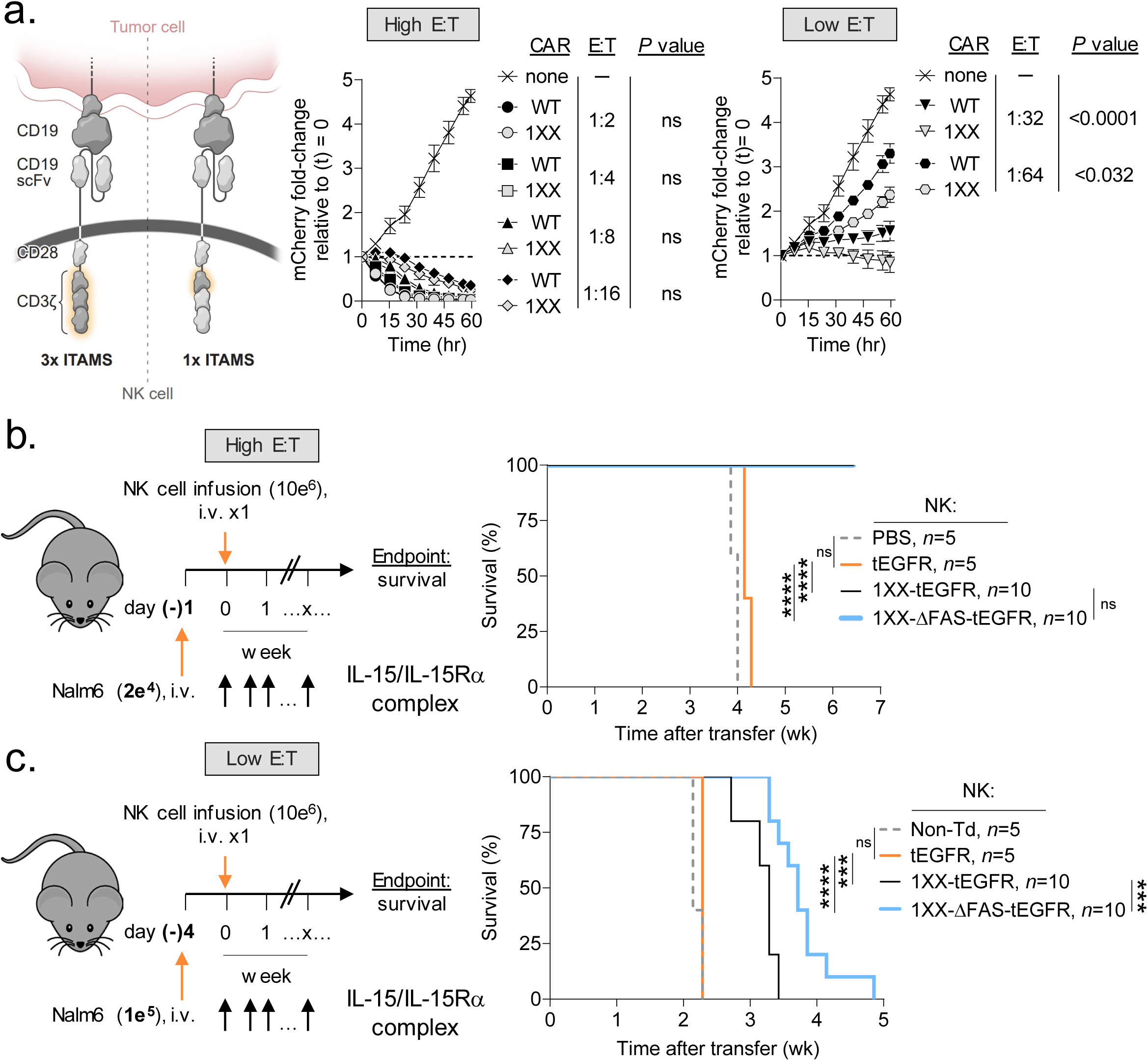
Disabling FAS-signaling enhances CAR-NK antitumor efficacy *in vivo*. (**a**) Comparison of the *in vitro* cytolytic efficiencies of NK cells transduced with a wild type (WT) or 1XX version of the 1928ζ CAR against Nalm6/mCherry at high versus low effector to target (E:T) ratios. Data shown as mean ± s.e.m. using *n*=3 biological replicates per condition. Statistical comparisons performed using a one-way ANOVA. ns = not significant, *P*>0.05. (**b**) Experimental design to compare the in vivo antitumor efficacy of human NK cells expressing the 1XX 1928ζ CAR ± a FAS dominant negative receptor (ΔFAS) against established Nalm6 B-ALL at a high versus low E:T ratio. All mice received a twice-weekly intraperitoneal injection of 1 μg of IL-15 pre-complexed with IL-15Rα-Fc (1:1 M). (**c**) Data plotted as a Kaplan–Meier survival curve (PBS, *n*=5; non-transduced NK cells, *n*=5; tEGFR alone, *n* = 5; 1XX 1928ζ-tEGFR, *n* = 10; 1XX 1928ζ-ΔFAS-tEGFR, *n* = 10). \*\*\*\**P*<0.0001, \*\*\**P*=0.001, and ns = not significant (*P*>0.05) using log-rank test. wk, week.

Based on these findings, we incorporated ΔFAS into a multi-cistronic vector encoding the 1XX CAR. We then compared the *in vivo* antitumor efficacy of a single IV infusion of 1XX CAR-NK cells that either express or do not express ΔFAS in mice bearing established Raji B-NHL at both a high and low E:T ratio. To control for potential intrinsic antitumor activity of NK cells, cohorts of mice received NK cells transduced with tEGFR alone. At a high E:T ratio, mice receiving 1XX CAR-NK cells either with or without ΔFAS co-expression significantly extended animal survival compared with NK cells transduced with tEGFR alone (**Fig. 6b**). By contrast, when cells were administered at a low E:T ratio as a stress test, we found that ΔFAS-expressing CAR-NK cells significantly prolonged survival compared with CAR-NK cells alone (**Fig. 6c**). Taken together, we conclude that co-expression of ΔFAS can enhance the *in vivo* antitumor potency of CAR-modified NK cells.

## Discussion

Herein, we demonstrate that FAS-L performs dichotomous functions in the context of CAR-T and CAR-NK therapies targeting B-cell malignancies (**Extended Data Fig. 5**). Whereas FAS-L limits CAR-modified lymphocyte persistence, it is expendable for CAR-mediated tumor lysis. Through construction and analysis of a single-cell transcriptomic atlas comprised of diverse cancer types and samples from patients who received CAR-T cells, we revealed that *FASLG* expression is highly restricted in humans. Rather than being expressed preferentially by malignant and stromal cells, as previously postulated^65,66^, we found that the dominant source of *FASLG* is endogenous T cells, NK cells, and CAR-modified lymphocytes. Relative to endogenous T cells, CAR-T cells displayed higher levels of *FASLG*, a finding attributable to recent encounter with antigen-bearing target cells. We further demonstrated that cellular activation promotes protein level expression of FAS-L in both CAR-T and CAR-NK cells.

Disabling FAS-signaling in CAR-expressing lymphocytes provided a fitness advantage following serial antigen-encounter *in vitro* and in competitive repopulation assays *in vivo*. These findings complement prior studies that found CAR designs which drive potent *in vitro* effector functions can lead to FAS-dependent T cell death and reduced persistence *in vivo*^67,68^. We found that the benefit of disrupting FAS extends beyond CAR-T to include CAR-NK cells, a lymphocyte population with favorable GVHD risk and cytokine release profiles but relatively low engraftment potential^60,69^. Augmented persistence resulting from cell-intrinsic blockade of FAS-signaling led to enhanced antitumor efficacy in multiple B-cell tumor models. Genetic ablation of *FASLG* in CAR-modified cells removed the benefit of FAS antagonism on cell survival. Thus, CAR-modified lymphocyte persistence is negatively self-regulated through a FAS-L:FAS interaction.

Unlike CAR-engineered lymphocyte survival, we established that on-target control of CD19^+^ tumors by CAR-T and CAR-NK cells is *FASLG* independent. We further demonstrated that while B-ALL and B-NHL are insensitive to recombinant FAS-L, both T and NK cells rapidly undergo apoptosis when exposed to the ligand. These findings parallel prior studies which reported that many leukemias and lymphomas display resistance to FAS cross-linking antibodies^70,71^. Correspondingly, we discovered that target cell expression of FAS is similarly dispensable for the antitumor activity of a 1928ζ CAR. This finding was confirmed by and extended to a second CAR containing a 41BB costimulatory domain using results from two CRISPR screens^56,57^. Mutations in *FAS* are observed in ∼5%-20% of B-cell malignancies^72–75^. However, these mutations frequently are subclonal^74^, suggesting they do not function as an escape mechanism from attack by endogenous immune cells. In the context of CAR-T cells, two recent studies performed next-generation sequencing (NGS) on B-NHL tumor DNA before and after cell infusion to identify tumor-intrinsic resistance mechanisms. In one study, which performed NGS on tumor biopsy samples, *FAS* mutations were not associated with treatment outcomes^76^. In a second study, circulating tumor DNA was analyzed prior to CAR-T cell infusion and at the time of disease progression^77^. Here also, mutations in *FAS* did not emerge under CAR-T cell selection *in vivo*. It is important to note that these findings do not exclude the possibility of FAS-dependent bystander killing by CAR-T cells, a phenomenon which may limit antigen-negative tumor escape^78,79^.

Beyond FAS, four additional members of the TNF death receptor family share dependency on the FADD adaptor protein, including TNF-R1, DR3, TRAIL-R1 (DR4), and TRAIL-R2 (DR5)^30,80^. *FADD* was identified as the gene most significantly associated with tumor resistance under selection by anti-CD19 CARs in two genome-scale CRISPR screens^56,57^. Both screens discovered that KO of *TNFSR10B*, the gene encoding TRAIL-R2/DR5, drove CAR-T resistance whereas *FAS* KO had no significant impact. Consistent with these findings, we identified that *TRAIL* KO significantly abrogated antitumor efficacy of anti-CD19 CAR-T cells, a finding corroborated by other investigators^56^. Of note, TRAIL has recently also been shown to be an essential effector mechanism for CAR-T targeting of a solid malignancy^81^. Together, these data support the conclusion that death receptor-signaling is an important determinant of CAR-mediated control of B-cell malignancies even if FAS-L/FAS interactions are non-essential. A limitation of our study is that it focused on B-ALL and B-NHL cancers. Establishing whether FAS-signaling is dispensable in other malignancies, particularly solid cancers, or using alternative therapies, such as bispecific engagers, remain important areas of future research.

In conclusion, our findings demonstrate that disruption of FAS-signaling is a broadly applicable strategy to enhance the persistence of genetically-redirected lymphocytes. Based on the observation that *FASLG* expression is restricted primarily to immune cells and not cancer or stromal cells, this approach should have utility across cancer types. While our studies focused on T and NK cells, a similar strategy may also benefit therapies employing invariant NK-T and γδ T cells as these lymphocyte subsets express and are responsive to FAS^82,83^. Although we employed a dominant negative receptor, cell-intrinsic FAS antagonism can be accomplished using alternative methods. These include synthetic switch receptors^27,28,33,34^, inhibitory RNAs^35^, and CRISPR-based genome editing^26,84^. Clinical trials (<u>NCT05617755</u>, <u>NCT06105021</u>) testing several of these strategies have recently been initiated and will provide additional evidence for whether FAS-L limits cellular persistence in humans.

## Extended data figure legends

**Extended Data Fig. 1:**
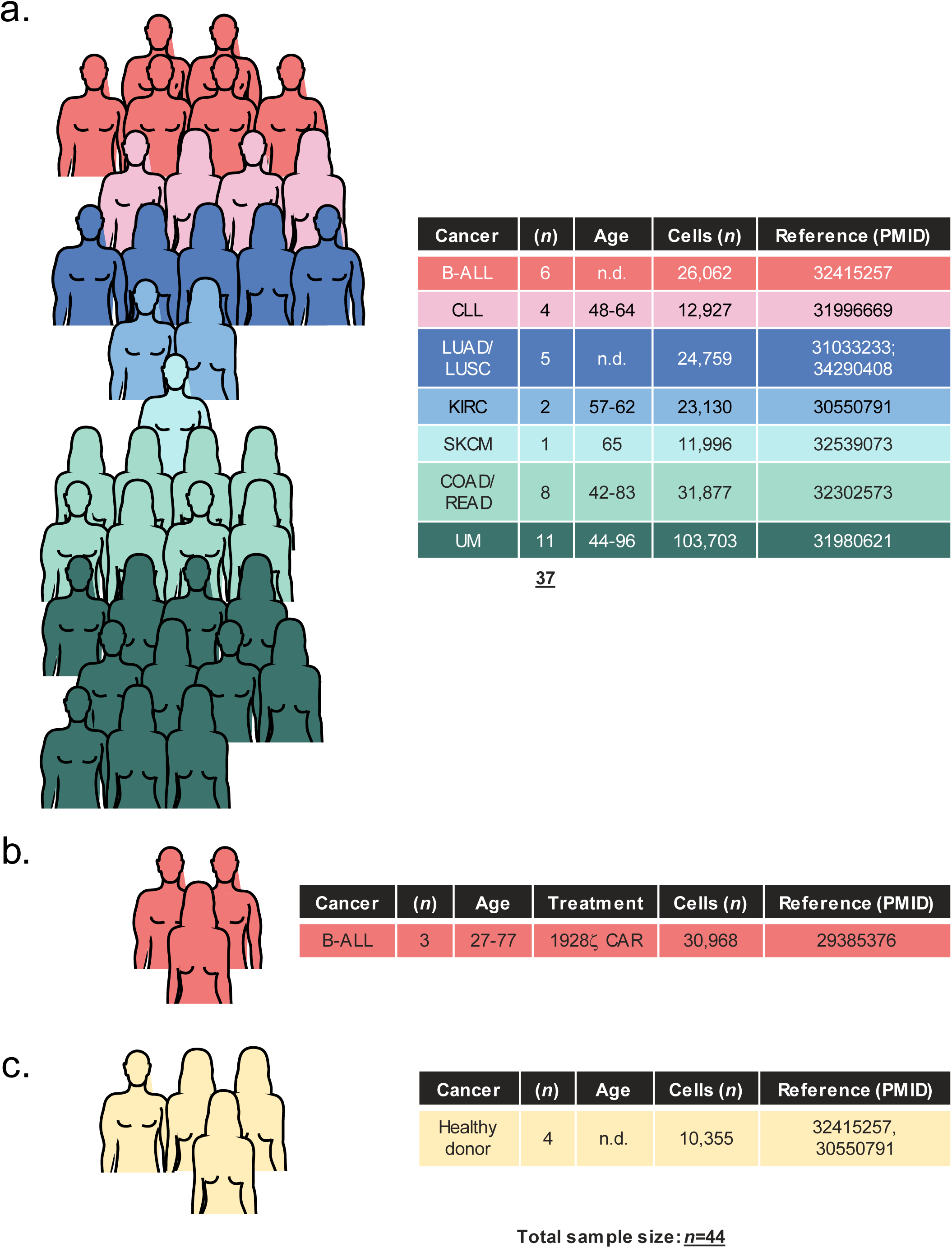
Overview of patient samples used to generate a single-cell atlas of *FASLG* expression by endogenous and CAR-engineered cells. Summary of (**a**) cell-therapy naïve cancer patients, (**b**) patients treated with a 1928ζ CAR, and (**c**) healthy donors analyzed using single-cell techniques to generate a *FASLG* expression atlas. Pictographs and tables display the sample size for each cancer cohort, gender distribution, age range, total number of single-cells analyzed, and reference to the primary data sets. B-ALL = B-cell acute lymphoblastic leukemia; CLL = chronic lymphocytic leukemia; COAD = colon adenocarcinoma; KIRC = kidney renal clear cell carcinoma; LUAD = lung adenocarcinoma; LUSC = lung squamous cell carcinoma; READ = rectal adenocarcinoma; SKCM = skin cutaneous melanoma; UM = uveal melanoma. n.d. = no data.

**Extended Data Fig. 2:**
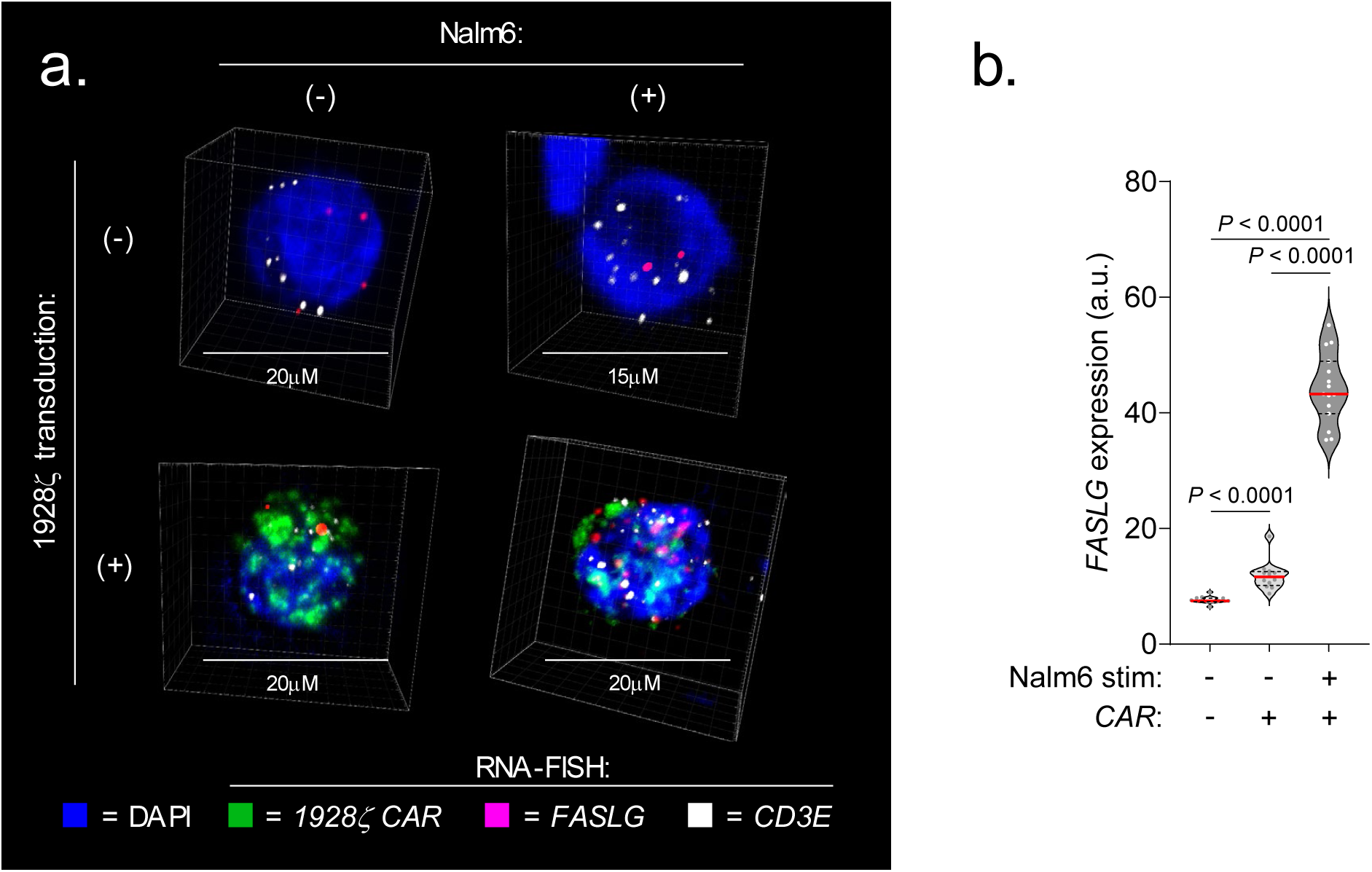
Quantification of *FASLG* expression by 1928ζ CAR-expressing T cells following *in vitro* co-culture with CD19^+^ leukemia cells using RNA *in situ* hybridization. (a) Representative immunofluorescent confocal images and (b) summary violin plots quantifying *FASLG* mRNA expression by non-transduced or 1928ζ CAR-expressing T cells at rest or 24h following in vitro co-culture with CD19^+^ Nalm6 B-ALL. Samples were co-hybridized with DAPI (blue) and multiplexed RNA-FISH probes specific for the mRNA sequence of the CAR’s single-chain variable fragment (scFv) (green), *CD3E* mRNA (white), and *FASLG* mRNA (red). Data shown is representative of results using T cells derived from *n*=2 healthy donors. Violin distributions are centered around the median (red horizontal line) with quartiles ranges displayed above and below (dashed horizontal lines). The maxima and minima are represented by the top and bottom of each plot. Each dot represents mean *FASLG* mRNA expression within a particular cell type from a region of interest. *P*-values calculated using a two-sided Student’s t-test. a.u. = arbitrary fluorescence units.

**Extended Data Fig. 3:**
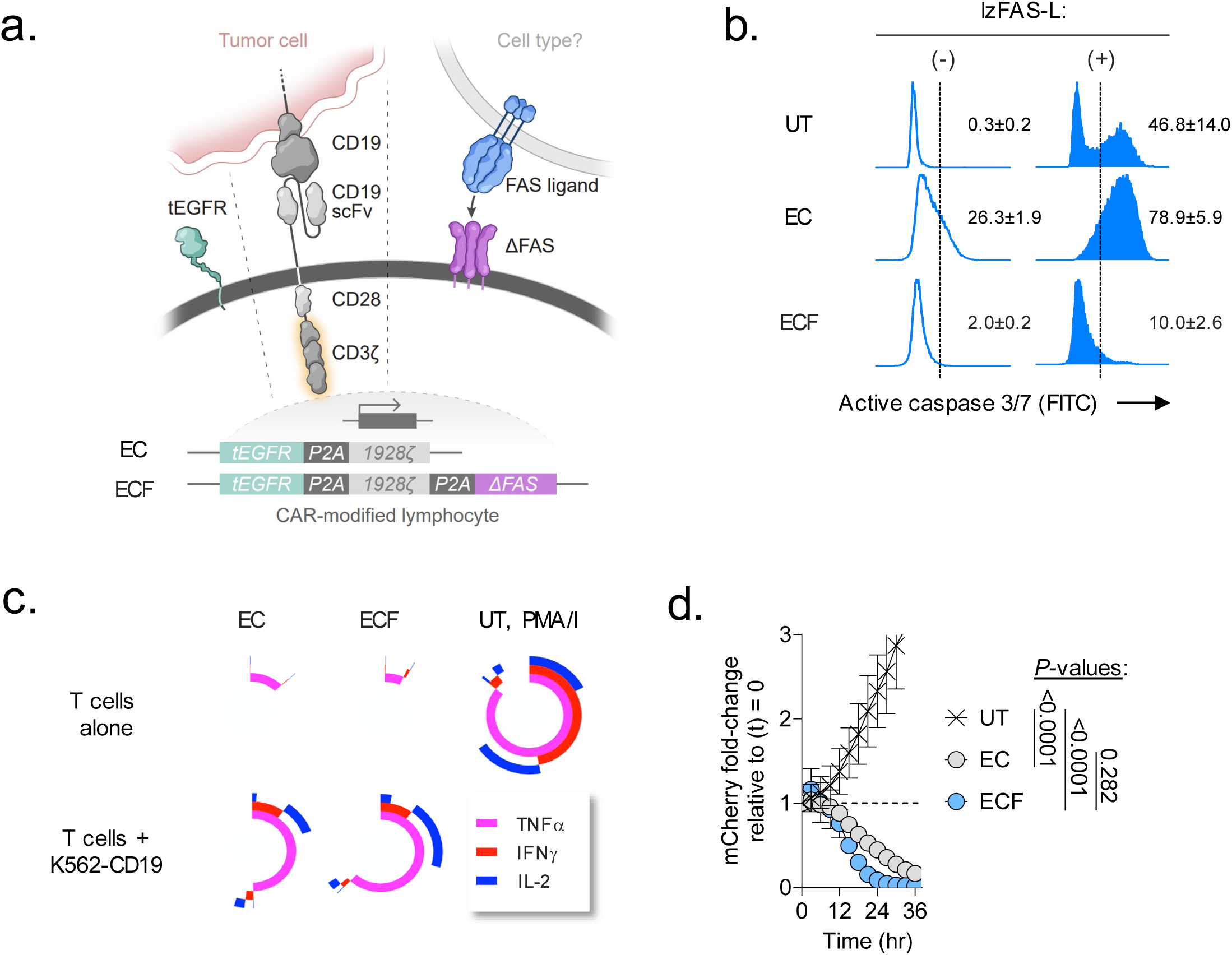
Cell-intrinsic disruption of FAS-signaling blocks CAR-T cell apoptosis but not effector function *in vitro*. (**a**) Schematic representation of two multi-cistronic vectors encoding tEGFR and a 1928ζ CAR alone (EC) or together with a FAS dominant negative receptor (ΔFAS) (ECF). To facilitate stoichiometric expression, each transgene was separated by a picornavirus self-cleaving 2A peptide sequence (P2A). To avoid vector recombination, each P2A in a single construct was encoded using degenerate codon sequence. (**b**) Representative FACS plots for activated caspase 3/7 in human T cells left untransduced (UT) as a control or transduced with an indicated multi-cistronic vector. Caspase activity was measured at rest and 4h following stimulation with 100ng mL^-1^ of a recombinant human FAS-L molecule oligomerized through a leucine zipper domain (lzFAS-L). Median of *n*=3 biologic replicates is shown together with mean ± s.e.m. of gated activate caspase 3/7^+^ lymphocytes. (**c**) Simplified Presentation of Incredibly Complex Evaluations (SPICE) analysis representing cytokine polyfunctionality of T cells transduced with indicated CAR constructs and co-cultured with or without K562-CD19 leukemia cells. T cells exposed to phorbol 12-myristate 13-acetate/ionomycin (PMA/I) was used as a positive control. Concentric plots indicate the median expression of three measured cytokines (TNFα, IFNγ, and IL-2) from *n*=3 biologic replicates. (**d**) *In vitro* cytolytic activity of UT, EC, or ECF-expressing T cells against Nalm6/mCherry. Data shown as mean ± s.e.m. using *n*=3 biological replicates per condition. *P*-values calculated using a one-way ANOVA.

**Extended Data Fig. 4:**
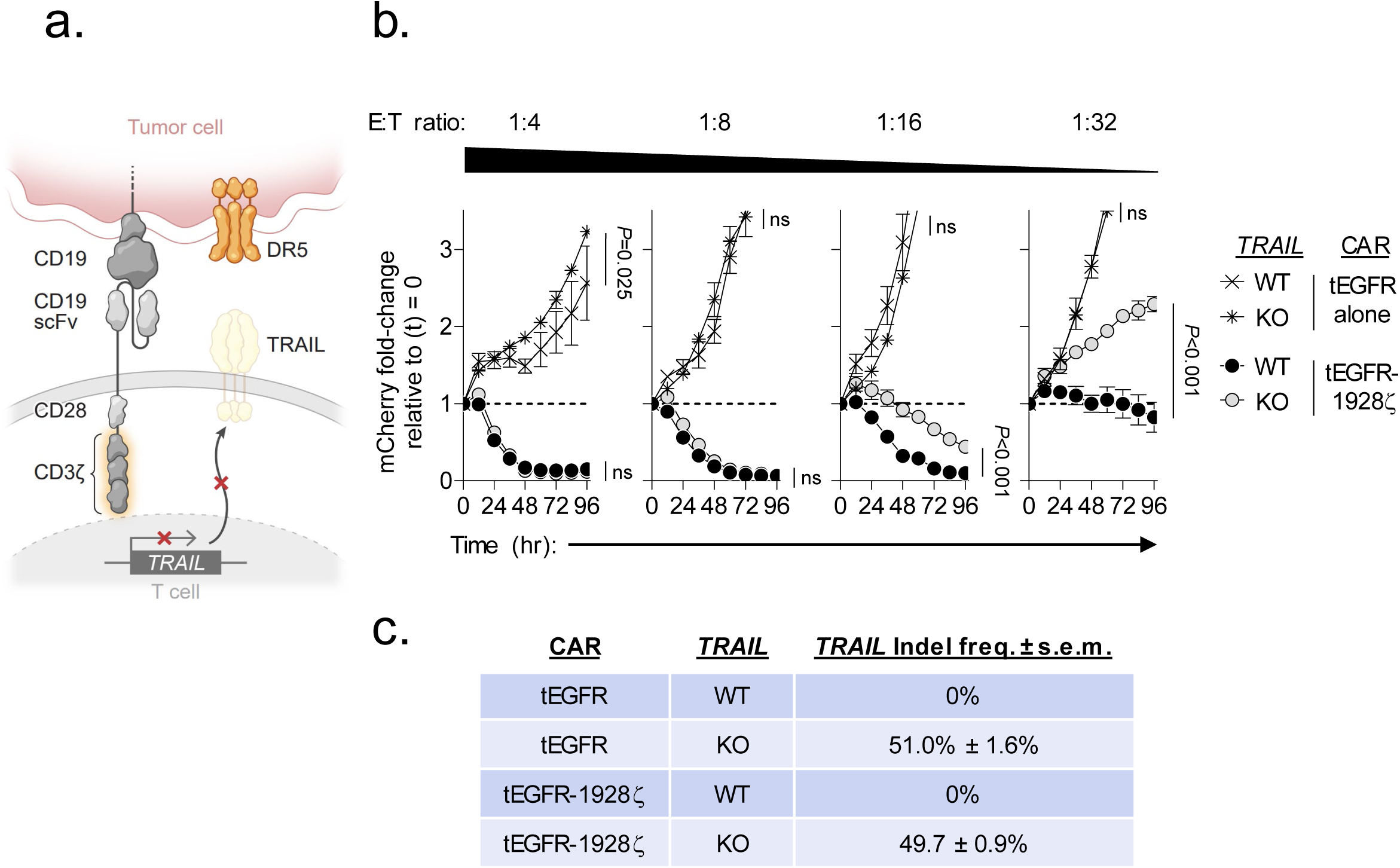
Knockout of *TRAIL* in 1928ζ CAR-T cells significantly impairs antitumor cytolytic activity. (**a**) Schematic for the CRISPR/Cas9-mediated knockout (KO) of *TRAIL* in human CD8^+^ T cells expressing a 1928ζ CAR. (**b**) Cytolytic activity of *TRAIL*-KO versus wild type (WT)-TRAIL 1928ζ CAR-T cells or T cells transduced with tEGFR alone against Nalm6/mCherry at indicated effector to target (E:T) ratios. Data shown as mean ± s.e.m. using *n*=3 biological replicates per condition. Statistical comparisons performed using a one-way ANOVA. ns = not significant, *P*>0.05. (**c**) Table displaying the measured frameshift insertion-deletion (Indel) frequency in *TRAIL* for each T cell group used in the cytolytic assay. Data shown as mean ± s.e.m. from *n*=3 technical replicates.

**Extended Data Fig. 5:**
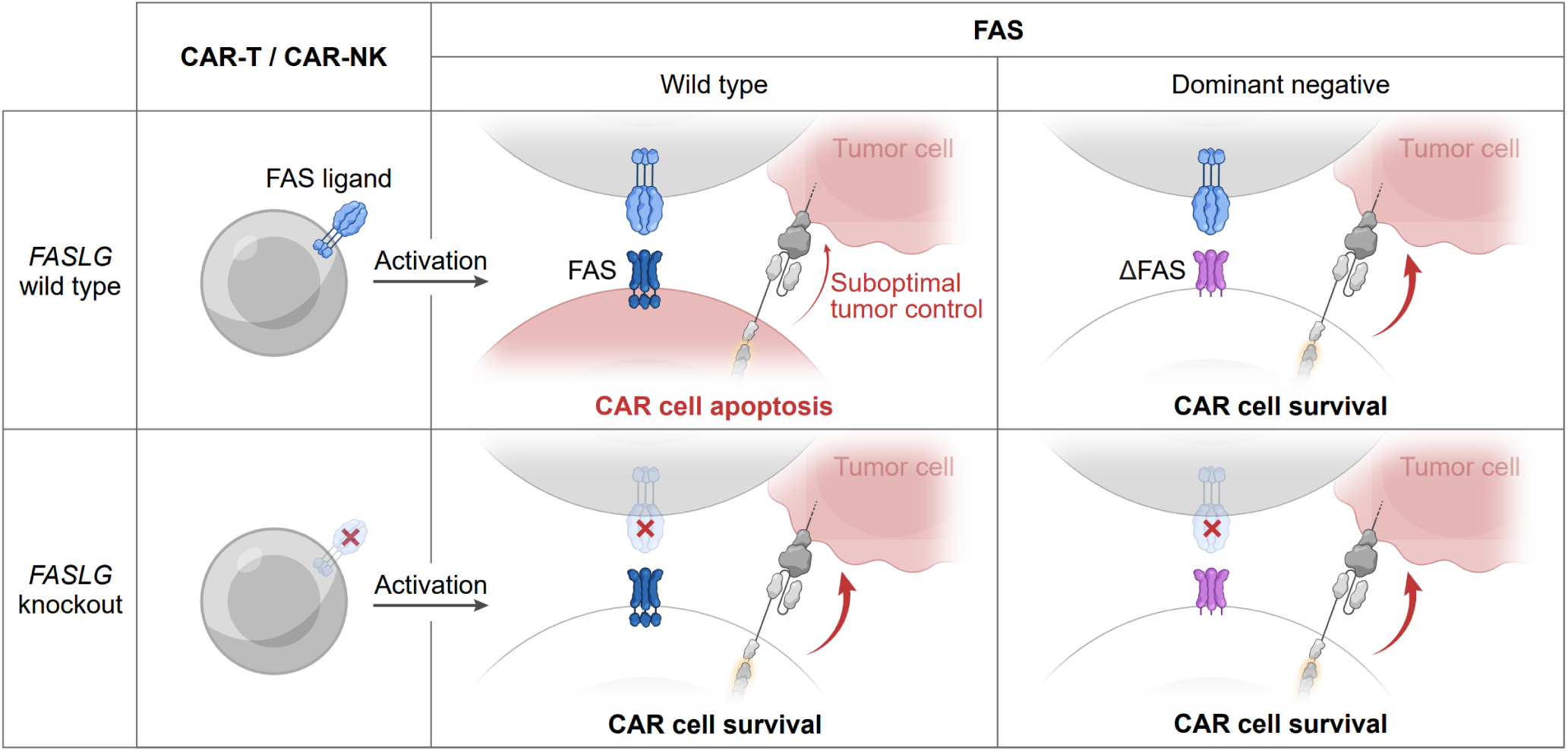
Model for the dichotomous functions of CAR-derived FAS-L on cellular persistence and antitumor efficacy. Cells colored in red indicate FAS-L induced apoptosis.

## METHODS

### Human materials and ethical approvals

Bone marrow aspirate samples were obtained from patients with relapsed B-ALL who received autologous 1928ζ CAR-T therapy between 2012 to 2013 on the Phase I MSKCC Institutional Review Board (IRB) approved protocol #09-114 (ClinicalTrials.gov ID: 01044069). As part of this study, all patients signed informed consent to collect biological specimens for research purposes. Bone marrow aspirates were collected in EDTA and bone marrow mononuclear cells (BMMC) were isolated on the day of procurement. Briefly, aspirates were centrifuged at 800*g* for 20 min at low acceleration and no deceleration, followed by aspiration of supernatant to a volume of 6 mL. Cells were resuspended in 4 mL of 2% of human serum antibody (HSA) PBS and the suspension was centrifuged again using the same settings. The buffy coat was extracted and washed once with 2% HSA PBS, and then once with 10 mL X-Vivo. Finally, the BMMC pellet was resuspended in 1 mL of 10% DMSO in HSA cryopreservation medium and stored at -80°C until transfer to liquid nitrogen.

### RNA-FISH staining and confocal imaging

BMMC were thawed and washed once with 1X PBS. Fresh cell pellets were placed on Superfrost™ Plus microscope slides (4951PLUS-001, Thermo Scientific), air dried for 20 min, fixed in 4% paraformaldehyde (PFA) for 15 min, and then washed 3x in PBS. Slides were dehydrated before being loaded into Leica Bond RX. Sections were pre-treated with EDTA-based epitope retrieval ER2 solution (Leica, AR9640) for 5 min at 95°C. The three experimental RNAscope™ FISH probes included: 1) synCD19scfv-C1 (ACD Bio, cat#1227848-C1) custom designed against mRNA for the single-chain variable fragment (scFv) of the 1928ζ CAR used on IRB#09-114, 2) Hs-FASLG-C2 (ACD Bio, cat#449058-C2) targeting human *FASLG* mRNA, and 3) Hs-CD3E-C3 (ACD Bio, cat#553978-C3) against human *CD3E* mRNA. Probes were hybridized for 2 h at 42°C. hs PPIB (ACD, Cat# 313918) and DapB (ACD, Cat# 312038) probes were used as positive and negative controls, respectively. The hybridized probes were detected using RNAscope™ LS Multiplex Reagent Kit (ACD Bio, Cat# 322800) according to manufacturer’s instructions. Alexa Fluor 488 tyramide (Life Technologies, B40953), CF 594 Tyramid (Biotium, 92174), and Alexa Fluor 647 tyramide (Life Technologies, B40958) were incubated with samples for 20 min at room temperature for fluorescence detection. After staining, slides were washed in PBS and incubated in 5 μg mL^-1^ 4’,6-diamidino-2-phenylindole (DAPI) (Sigma Aldrich) in PBS (Sigma Aldrich) for 5 min. PBS was rinsed, and slides were mounted in Mowiol 4-88 (Calbiochem). Slides were scanned and stored at -20°C prior to confocal imaging. Slides were annotated on QuPath and gated for DAPI, Alexa Fluor 488 (scFv of 1928ζ) and presence or absence of Alexa Fluor 647 (*CD3E*) positivity. The fluorescence intensity of CF 488 (*FASLG*) within the nucleus (DAPI) was measured for each cell and represented as arbitrary units. This value was then averaged over all cells meeting the gated expression criteria for each annotated region. For 3D image reconstructions, cells were imaged using the Zeiss LSM 880 confocal microscope. Z-stacks were generated with Zen lite and 3D images were extrapolated with Imaris imaging software.

### Cell lines and tissue culture

293GP cells (Takara Bio, 631458), Nalm6 (ATCC, CRL-3273), and Raji (ATCC, CCL-86) were purchased from commercial vendors. Nalm6-GFP/Luc cells were previously described^50^ and obtained from the Sadelain lab (MSKCC). Raji-GFP/Luc cells were also previously described^85^ and obtained from the Brentjen’s lab (MSKCC). K562-CD19 cells were previously described^86^ and provided under MTA from the Feldman lab (NIH). K562 Clone 9 cells were previously described^87^ and provided under MTA from the Lee lab (Nationwide Children’s Hospital). Raji and Nalm6 cell lines with stable NLS-mCherry or NLS-GFP reporter expression were developed by retroviral transduction. K562-CD19 *FASLG*-KO cells were generated by CRISPR gene editing, described below, followed by single cell cloning. All tumor cell lines were cultured in RPMI 1640 (Gibco, 11875085) supplemented with 0.5% v/v Penstrep (Gibco, 15070-063), 1% v/v HEPES (Gibco, 15630-080), 0.02% v/v Gentamycin (MP Biomedicals, 1676245) and 10% v/v heat inactivated fetal bovine serum (GeminiBio, 100-106).

### Plasmids design, viral packaging, and transduction

The 1928ζ and 1XX 1928ζ sequences were previously reported^50,64^ and provided by the Sadelain lab (MSKCC). 1928ζ, tEGFR and ΔFAS sequences were separated by P2A motifs and synthesized by Genscript. The synthesized genes were subcloned into a SFG^88^ retroviral vector and the identity of the new sequences was confirmed by Sanger sequencing (Genscript). Similarly, NLS-mCherry and NLS-GFP sequences were synthesized and cloned into an SFG vector by Genscript. For transfection of the plasmids mentioned above, 293GP cells were co-transfected with 4.5 μg pMSGV-1 and 2.25 μg RD114 feline envelop plasmid provided under MTA from the Rosenberg Lab (NIH) using Lipofectamine™ 3000 Transfection Reagent (Thermo Fisher, L3000075). Viral supernatant was harvested on day 2 post transfection and used fresh. For viral transduction, T cells were activated for 2 days and NK cells were activated for 4 days followed by retronectin (Takara, T100B) mediated viral transduction. In brief, retronectin was coated on non-tissue culture plates at 10 μg mL^-1^ in PBS, 500 μL per well in non-tissue culture 24-well plates overnight. On the day of transduction, coated plates were washed by PBS and blocked by 2% BSA in PBS for 30 min at room temperature. Viral supernatant was loaded on aspirated plate and spun at 2500*g* for 2 h at 32°C. Viral supernatant was aspirated after centrifugation. Cells were loaded onto plates and spun at 300*g* for 30 min at 32°C. Transduction were confirmed by flow cytometry on day 2 (for T cells and cell lines) or day 5 (for NK cells). The E:T ratio used for both *in vitro* and *in vivo* experiments was calculated based on transduction rates determined using tEGFR^+^ or LNGFR^+^ lymphocytes, as assessed by flow cytometry data.

### Primary cell preparation

For T cell preparations, buffy coats were obtained from the New York Blood Center or, alternatively, Leukocyte Reduction System cones (STEMCELL Technologies, 200-0093). T cells were isolated by EasySep™ Human T Cell Isolation Kits (STEMCELL Technologies, 17951). Naïve T cells were isolated by EasySep™ Human Naïve Pan T Cell Isolation Kits (STEMCELL Technologies, 17961). Naïve CD8^+^ T cells were isolated by EasySep™ Human Naïve CD8^+^ T Cell Isolation Kit (STEMCELL Technologies, 19258). All isolated T cells were activated by CD3/CD28 Dynabeads (Thermo Fisher, 11132D) at a 1:1 ratio for 2 days. For NK cell preparations, cord blood mononuclear cells were ordered from STEMCELL Technologies (STEMCELL Technologies, 70007.1 or 70007.2). NK cells were isolated by EasySep™ Human NK Cell Isolation Kits (STEMCELL Technologies, 17955) and activated by 100 Gy irradiated K562 Clone 9 feeder cells. Feeder cells were mixed with NK cells at a 2:1 tumor:NK cell ratio. Primary cells were cultured in RPMI 1640 (Gibco, 11875085) with 1% v/v Penstrep (Gibco, 15070-063), 2.5% v/v HEPES (Gibco, 15630-080), 0.02% v/v Gentamycin (MP Biomedicals, 1676245) and 10% v/v heat inactivated human serum AB (GeminiBio, 100-512). Recombinant human IL-2 (PeproTech, 200-02) was added to T cells and NK cells cultures at 50 IU mL^-1^ or 200 IU mL^-1^, respectively.

### Flow cytometry

For surface staining, cells were washed with PBS and stained with antibodies at 4°C for 30 min in PBS supplemented with 0.5% FBS. Cells were washed twice with PBS + 0.5% FBS and acquired on an LSRFortessa X-20 (BD). Antibodies used for surface staining were derived from the following clones: AffiniPure Fab Fragment Goat Anti-Mouse IgG (polyclonal, Jackson ImmunoResearch) CCR7 (G043H7, Biolegend), EGFR (AY13, Biolegend), LNGFR (C40-1457, BD), FAS (DX2, Biolegend), CD3 (OKT3, Biolegend), CD4 (SK3, Thermo Fisher), CD8a (RPA-T8, Biolegend), CD45 (H130, Biolegend), CD45RA (HI100, Biolegend), CD56 (NCAM, Biolegend), PD-1 (MIH4, BD). LIVE/DEAD Fixable Aqua Dead Cell Stain Kit (Thermo Fisher) was used for live cell gating in all flow cytometry experiments. For NK cells, Human TrueStain FcX blocking solution (Biolegend, 422302) was added to cells for 10 min before staining. Cells were washed twice with PBS + 0.5% FBS before acquisition or used for further staining. For intracellular staining, T cells or NK cells were mixed with tumor cells for 6 h in the presence of GolgiPlug (BD, BDB555029). As a positive control, eBioscience™ Cell Stimulation Cocktail (500X) (Thermo Fisher, 00-4970-03) was added during the same 6 h incubation. After staining using surface antibodies and LIVE/DEAD, cells were permeabilized and fixed by Fixation/Permeablization Kit (BD, 555028) for 20 min at 4°C. Cells were then washed once in Perm/Wash buffer (BD, 51-2091KZ) and stained by anti-IFNγ (BD, B27), anti-TNFα (Thermo Fisher, Mab11), and anti-IL2 (Thermo Fisher, MQ1-17H12) for 40 min at 4°C. Cells were washed twice with Perm/Wash buffer before acquisition. Details about the gating strategy applied to each panel in the flow cytometry immunophenotyping analysis are described in **Supplementary Fig. 1**.

### Apoptosis assay

lzFAS-L was made from transfected HEK293T cells, as previously described^54^. Cells were treated with lzFAS-L at designated time points. For active Caspase-3/7 staining, CellEvent™ Caspase-3/7 Green Detection Reagent (Thermo Fisher, C10423) was added to cultures at 20 μM and incubated for 30 min at 37°C. Cells were washed twice with Annexin V Binding Buffer (Biolegend, 422201) and stained with APC Annexin V (Biolegend, 640941) at 5% v/v in the presence of antibodies for other surface markers. Cells were washed twice with Annexin V Binding Buffer before acquisition by FACS.

### CRISPR gene editing

sgRNA (Synthego, 50 μM) and NLS-Cas9 protein (Synthego, 20μM) were mixed at 2.5:1 molar ratio (1 μL sgRNA + 1 μL protein) for 10 min at room temperature. 1 x 10^6^ cells were resuspended with 18 μL electroporation buffer, P3 buffer (Lonza, V4XP-3032) for primary cells, SG buffer (Lonza, V4XC-3032) for Raji cells, and SF buffer (Lonza, V4XC-2032) for Nalm6 cells and K562 cells. 2 μL of Cas9/sgRNA complex was added to 18 μL of cells. Cells were electroporated on a Lonza 4D-Nucleofector with X Unit using vendor’s recommended settings for each cell type. Cells were transferred to flasks with warmed media immediately after electroporation. To verify gene editing, DNA was extracted from CRISPR edited cells using DNsasy Blood & Tissue Kit (QIAGEN, 69506). DNA samples were PCR amplified using KOD Hot Start DNA Polymerase (MilliporeSigma, 71842) with the primers listed in **Supplementary Table 1** and the PCR product was subjected to Sanger sequencing. Indel frequencies were quantified using the ICE CRISPR Analysis Tool (Synthego).

### Western blot

To determine FAS-L protein expression, 5 x 10^6^ to 1 x 10^7^ cells per sample were lysed by RIPA buffer (Thermo Fisher, 89900) and shaken on a rocker for 1 h at 4°C. Cell lysates were further sonicated on Bioruptor® Pico (diagenode) for 20 cycles (30 s sonication, 30 s cool-down) in ice water. Lysates were added with LDS buffer, protease inhibitor, and sample reducing buffer (Thermo Fisher, NP0004) and loaded on Mini-PROTEAN TGX gel (Bio-Rad, 4561094). Precision Plus Protein™ was loaded to the gel as molecular weight standard. Gels were run at 100V until the dye ran out of the gel. Gel was transferred on a PVDF membrane using Transblot Turbo system (Bio-Rad). The membrane was washed in deionized water 3 times for 5 min, blocked by 5% nonfat milk in TBST (Bio-rad) for 30 to 60 min at room temperature with agitation, and stained by Rabbit anti-human FAS-L (Cell Signaling, D1N5E) at 1:500 v/v. The membrane was rocked overnight at 4°C with light avoided then washed 3 times with agitation for 5 min and stained by secondary antibody IRDye 800 CW Goat anti Rabbit (LI-COR, 926-32211) at 1:20,000 v/v. Lastly, the Membrane was scanned on an Odyssey CLx system (LI-COR). For housekeeping protein blotting, the same membrane was soaked in NewBlot PVDF stripping buffer (LI-COR, 928-40032) for 15 min followed by the same staining steps as described above using primary antibody β-Actin (Cell Signaling, 8H10D10) at 1:1,000 v/v and secondary antibody IRDye 680RD Goat anti Mouse (LI-COR, 926-68070) at 1:20,000 v/v.

### *In vitro* lymphocyte co-culture restimulation assays

1 x 10^6^ tLNGFR-1928ζ and 1 x 10^6^ tEGFR-1928ζ-ΔFAS expressing cells were mixed at 1:1 ratio on day 0. For CAR-T cells, 1 x 10^6^ K562-CD19 *FASLG* KO tumor cells were added to co-culture on day 0, 9 and 14. Cells were stained for flowcytometry on day 0, 2, 7, 9, 12, 14 and 17. For CAR-NK cells, 1 x 10^6^ K562 Clone9 *FASLG* KO tumor cells were added to co-culture on day 0, 5, 10 and 15. Cells were stained for FACS on day 0, 3, 5, 8, 10, 13, 15, 18 and 20. Cell culture media with fresh IL-2 were changed every 2 to 3 d during the experiments. Media volumes were measured and changed with fresh media at the same volumes.

### Real-time live cell imaging assays

For tumor cell killing assays, GFP-NLS or mCherry-NLS expressing tumor cells were loaded onto 96 well plates (Corning, 353072) at 1 x 10^5^ cells per 100 μL in complete media. Plates were left at room temperature for 20 min and transferred to the Incucyte SX1 instrument (Sartorius) for baseline images. The plate was then removed from the Incucyte and loaded with lymphocytes at designated ratios. The Plate was left at room temperature for 20 min again and transferred to the Incucyte for imaging. Analysis was done using Incucyte Basic Analysis Software. For lzFAS-L apoptosis assays, mCherry-NLS expressing tumor or T cells were loaded onto a 96 well plate at 2 x 10^5^ cells per 100 μL. Cells were treated with indicated concentrations of lzFAS-L. The same imaging and analysis procedures were used as for the tumor killing assays.

### ACT xenograft models

All mouse experiments were performed in accordance with an MSKCC Institutional Animal Care and Use Committee-approved protocol (19-08-013). Six to 10-week-old female NOD.Cg-Prkdcscid Il2rgtm1Wjl/SzJ (NSG) mice were used for all animal experiments (Jackson Laboratory) and housed in pathogen-free conditions at the MSKCC vivarium. The mouse room maintained a 12-hour light/dark cycle, temperature of 65–75 °F and humidity levels of 40–60%. Nalm6 GFP/Luc or Raji GFP/Luc cells were given by tail vein in 200μL PBS. CAR-T, CAR-NK, or control cells were also administered by tail vein injection in 200μL PBS 1 day or 4 days post tumor inoculation. For mice that received NK cells, IL-15/IL-15Rα complex was administered intraperitoneally twice weekly for the duration of the experiment, starting at the time of the first NK cell transfer. The complex was freshly prepared by incubating recombinant human IL-15 (Peprotech, 200-15) with IL-15Rα-Fc (R&D Systems, 7194-IR) at a 1:1 molar ratio for 30 min at 37 °C and administered at a final concentration of 1 μg of IL-15 per mouse. Tumor burden was measured by bioluminescence using IVIS Imaging System (PerkinElmer). Mice health conditions were monitored by MSKCC Research Animal Resource Center (RARC). Euthanasia decisions were determined based using pre-determined clinical criteria and was performed at the discretion of the RARC veterinary staff who were blinded to the identity of the treatment groups.

### Tissue collection and sample preparation

Tissues were dissected following CO_2_ euthanasia and kept in PBS with 1mM EDTA (EMD Millipore, 324504). Tissues were processed using 100 μm cell strainers (Falcon, 352360). Processed cells were centrifuged at 1200 rpm for 5 min and resuspended in ACK lysing buffer (Gibco, A10492) for 20 min at room temperature. Cells were filtered through Flowmi 70 μm cell strainers (Bel-Art, 136800070). Cells were washed once with PBS supplemented with 1mM EDTA and resuspend in FcR blocking buffer using Human TruStain FcX (Biolegend, 422302) and Mouse TruStain FcX PLUS (Biolegend, 156604) at room temperature for 15 mins. Cells were washed twice before proceeding with antibody staining.

### Single cell RNA sequencing datasets and processing

We collected publicly available single cell RNA sequence datasets from eight studies listed in **Extended Data Fig. 1**. Raw count matrices were retrieved from GEO (GSE132509, GSE111014, GSE117570, GSE111360, GSE148190, GSE146771, GSE176021, GSE139829) and cell annotations (when available) were retrieved from TISCH^89^. TISCH datasets were uniformly processed with MAESTRO^90^ (which included removal of low-quality cells, cell-type annotation, and malignant cell identification). Only TISCH annotated (thus filtered) cells were retained. For datasets GSE111360 and GSE176021 lacking TISCH annotations, cells were filtered out if the number of detected genes was <500, and features were removed if they were expressed in <10 cells; cell types were then annotated with SingleR v.1.0.6^91^ with reference to the TISCH-annotated datasets. Cells from all datasets were combined and integrated with the fastMNN function from batchelor v.1.10.0^92^ using the top 2,000 most variable genes. Dimensionality reduction was performed with runUMAP from scater v.1.22.0^93^ on the batch-corrected values. Size-factor normalized log-counts were obtained via computeSumFactors from scran v1.22.1^94^ and logNormCounts from scater v1.22.0. UMAPs and violin plots of *FAS*/*FASLG* were visualized with scater v1.22.0 and dittoSeq v1.6.0^95^. Code is available at https://github.com/abcwcm/Yi2024.

### Whole-genome CRISPR screen re-analysis

Count matrices from two whole genome screens were obtained either from supplementary tables^57^ or from Gene Expression Omnibus (GSE130663)^56^ and reanalyzed using MAGeCK (v0.5.9.2) with default parameters^96^.

### Statistics and reproducibility

No statistical methods were used to predetermine sample size. Appropriate statistical tests were used to analyze data, as described in each figure legend. Statistical analyses were performed with GraphPad Prism version 10 software. Significance was preset at *P*<0.05. *In vitro* experimental data were generated from two or more independent experiments containing *n*=3 biological replicates per condition per experiment. For *in vivo* mouse experiments, treatment groups had *n* = 10-15 mice per condition, and control groups had *n* = 5-10 mice. An investigator blinded to treatment groups performed both data acquisition and data analysis. *In vivo* experiments were independently repeated two times, and all attempts at replication were successful. For both *in vitro* and *in vivo* experiments, no data were excluded from any analysis.

## Acknowledgements

This study was supported in part by NIH R37 CA259177, P50 CA217694, and P30 CA008748 (C.A.K.); Mr. William H. Goodwin and Mrs. Alice Goodwin and the Commonwealth Foundation for Cancer Research (C.A.K.); The Center for Experimental Therapeutics at Memorial Sloan Kettering Cancer Center (C.A.K.); The MSK Technology Development Fund (C.A.K. and C.S.H.); The Parker Institute for Cancer Immunotherapy (C.A.K.); The Sarcoma Center at MSKCC (C.A.K.); The Damon Runyon Cancer Research Foundation CI-96-18 (C.A.K.); the Geoffrey Beene Cancer Research Center (C.A.K.); the Tow Center for Developmental Oncology (C.A.K.), and Cycle for Survival (C.A.K.). F.Y. was supported in part by an immuno-oncology research fellowship from the William Randolph Hearst Foundation and the Parker Institute for Cancer Immunotherapy. T.C. was supported in part by a Career Development Award from the Tow Center for Developmental Oncology. C.S.H. was supported in part by the MSK Paul Calabresi Career Development Award for Clinical Oncology (5K12 CA184746). K.N.K. was supported by the German Research Foundation (DFG) Fellowship. Original artwork featured in figures 4, 5, 6 and extended figures 3-6 was performed with the assistance of SciStories^®^.

## Author contributions

F.Y. and C.A.K. conceived of and designed all experiments. F.Y., T.C., N.Z., R.E., E.J.B., H.A., M.V.G., C.S.H., I.E., and S.S.C. performed or supported *in vitro* experiments. F.Y. and T.C. performed and analyzed RNA-FISH confocal imaging and Western blot experiments. F.Y., T.C., N.Z., and R.Z. generated cell products and analyzed data resulting from *in vivo* animal studies. F.D., P.Z., and D.B. assembled and analyzed scRNA-seq data using publicly available data. K.N.K. re-analyzed results from publicly available CRISPR screens. J.H.P. contributed clinically annotated patient samples. J.F., S.S.C., K.C.H., and M.S. provided key materials, reagents, and technical know-how. K.C.H. and M.S. provided additional study supervision. C.A.K. and F.Y. wrote the initial and revised manuscripts with input and contributions from all authors. All authors approved the revised manuscript.

---

**Supplementary Data Fig. 1:**
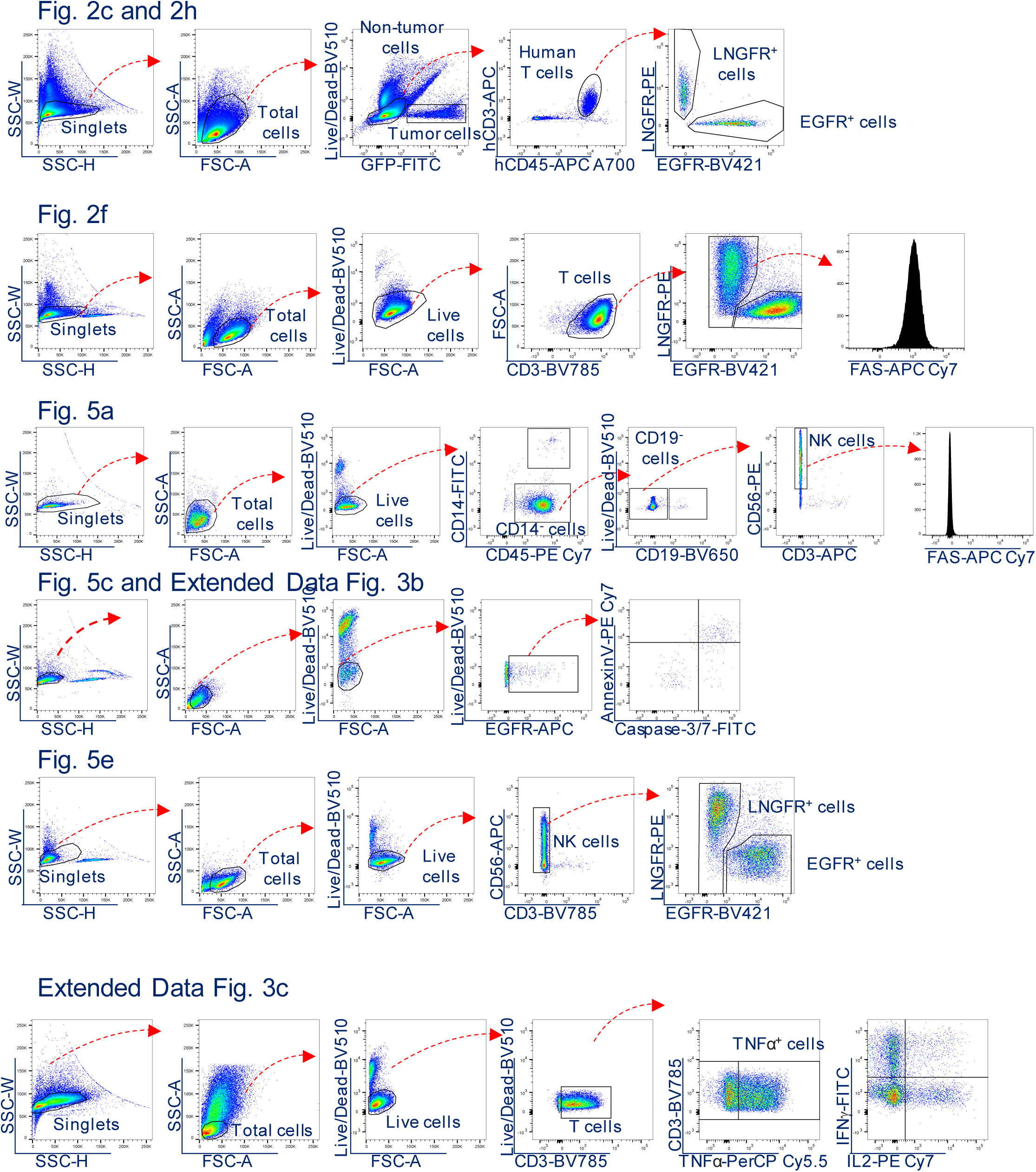
Gating strategy for the immunophenotyping by FACS analysis for CAR-T and CAR-NK cells.

**Supplementary Table 1:**
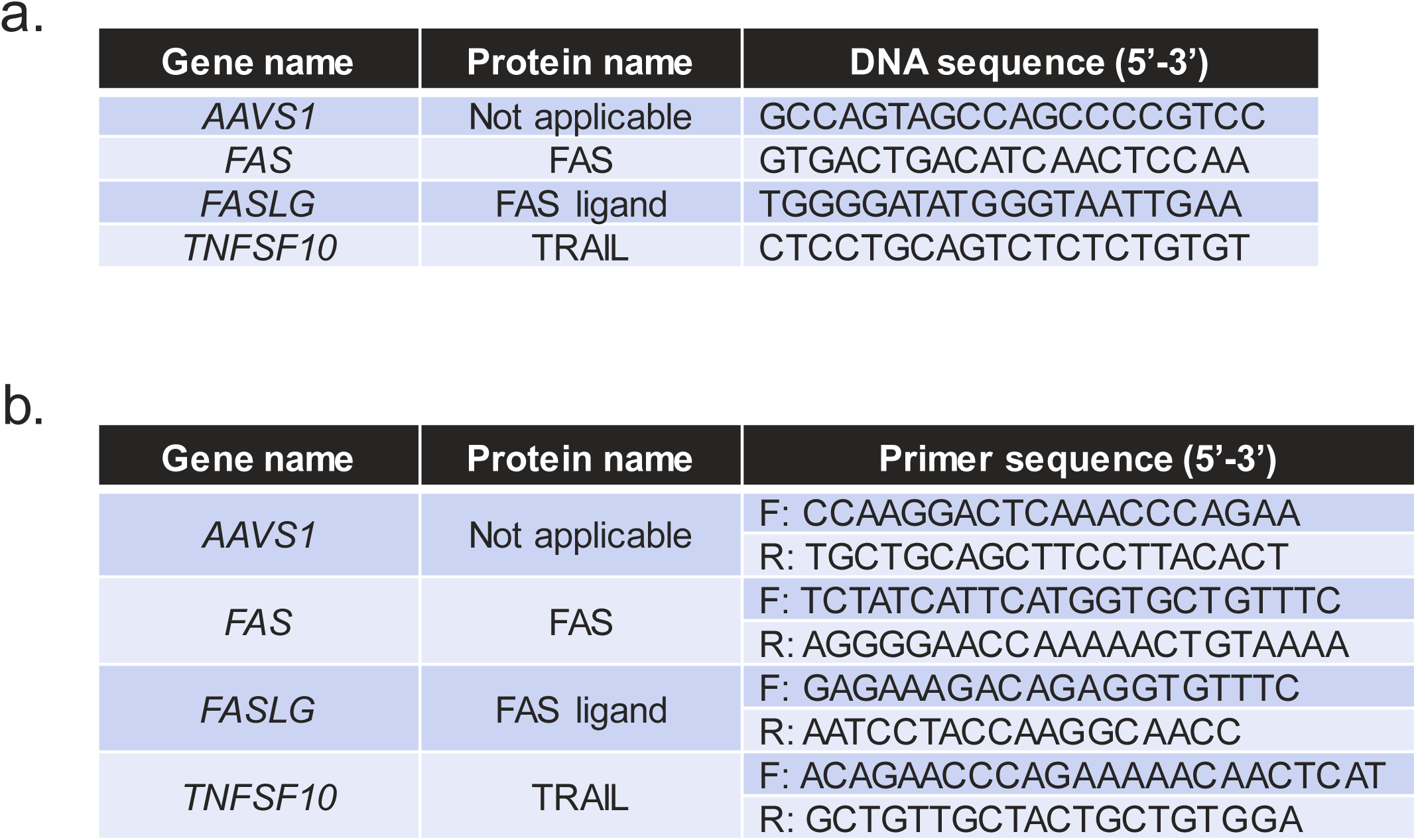
sgRNA sequences for CRISPR/Cas9 gene editing.

